# Kinesin-6 Klp9 orchestrates spindle elongation by regulating microtubule sliding and growth

**DOI:** 10.1101/2021.02.17.431708

**Authors:** Lara K. Krüger, Matthieu Gélin, Liang Ji, Carlos Kikuti, Anne Houdusse, Manuel Théry, Laurent Blanchoin, Phong T. Tran

## Abstract

Mitotic spindle function depends on the precise regulation of microtubule dynamics and microtubule sliding. Throughout mitosis, both processes have to be orchestrated to establish and maintain spindle stability. We show that during anaphase B spindle elongation in *S. pombe*, the sliding motor Klp9 (kinesin-6) also promotes microtubule growth *in vivo. In vitro*, Klp9 can enhance and dampen microtubule growth, depending on the tubulin concentration. This indicates that the motor is able to promote and block tubulin subunit incorporation into the microtubule lattice in order to set a well-defined microtubule growth velocity. Moreover, Klp9 recruitment to spindle microtubules is dependent on its dephosphorylation mediated by XMAP215/Dis1, a microtubule polymerase, to link the regulation of spindle length and spindle elongation velocity. Collectively, we unravel the mechanism of anaphase B, from Klp9 recruitment to the motors dual-function in regulating microtubule sliding and microtubule growth, allowing an inherent coordination of both processes.

## Introduction

Mitotic spindle assembly and function requires microtubules to undergo alternating periods of growth and shrinkage together with the action of molecular motors, that can crosslink and slide apart spindle microtubules. A fine balance of microtubule dynamics and microtubule sliding is essential to achieve faithful chromosome separation in all mitotic processes (Cande and Mcdonald, 1986; Cande and McDonald, 1985; Cheerambathur et al., 2007; Yukawa et al., 2019b). Yet, while work has mainly been focused on each process independently, it is unclear how they are coordinated.

During anaphase B, the spindle elongates to push apart the spindle poles and separate the two chromosome sets (Mallavarapu et al., 1999; Oegema et al., 2001; Roostalu et al., 2010; Scholey et al., 2016; Straight et al., 1998; Vukušić et al., 2017). While in the fungus *Ustilago maydis* and Ptk1 cells spindle elongation is mainly achieved through cortical pulling forces acting on astral microtubules (Aist et al., 1993; Fink et al., 2006; Grill et al., 2001), these forces are not present or dispensable in a plethora of other species. In yeast, *C. elegans, D. melanogaster*, plants, and human, spindle elongation is realized by the generation of microtubule sliding forces at the spindle midzone (Euteneuer et al., 1982; Khodjakov et al., 2004; Kiyomitsu and Cheeseman, 2013; Redemann et al., 2017; Sharp et al., 1999; Tolic-Norrelykke et al., 2004; Vukušić et al., 2019; Yu et al., 2019). The spindle midzone refers to the microtubule overlap at the spindle center, formed by antiparallel-oriented microtubules (interpolar microtubules), which originate from the two opposite spindle poles (Ding et al., 1993; Euteneuer et al., 1982; Mastronarde et al., 1993; Mcdonald et al., 1977; Mcintosh and Landis, 1971; Ward et al., 2014; Winey et al., 1995).

Microtubule sliding is promoted by members of the kinesin-5 family in most organisms (Avunie-Masala et al., 2011; Brust-Mascher et al., 2009; Saunders et al., 1995; Sharp et al., 2000; Straight et al., 1998). The tetrameric kinesin crosslinks antiparallel microtubules and slides them apart (Avunie-Masala et al., 2011; Brust-Mascher et al., 2009; Kapitein et al., 2005; Saunders et al., 1995; Shimamoto et al., 2015; Straight et al., 1998). By walking towards the microtubule plus-ends of each of the two microtubules, that it crosslinks, kinesin-5 can move the microtubules relative to each other (Kapitein et al., 2005). In fission yeast, this task is mainly performed by the kinesin-6 Klp9, which localizes to the spindle midzone upon anaphase onset (Fu et al., 2009; Krüger et al., 2019; Rincon et al., 2017; Yukawa et al., 2019a). The kinesin-5 Cut7 also generates microtubule sliding forces, but to a lower extent than Klp9 (Rincon et al., 2017). In absence of both sliding motors, spindle elongation is abolished, leading to ‘cut’ of chromosomes by the cytokinetic ring and mis-segregated chromosomes (Rincon et al., 2017).

Concomitantly with microtubule sliding, the interpolar microtubules have to grow at a rate that allows the spindle to elongate while maintaining the spindle midzone (Cande and Mcdonald, 1986; Cande and McDonald, 1985; Cheerambathur et al., 2007; Masuda, 1995; Masuda and Zacheus, 1987; Saxton and Mclntosh, 1987). Microtubule growth has to occur with at least the velocity the spindle microtubules slide apart to keep the microtubule overlap, necessary for spindle stability. Microtubule dynamics during anaphase B have been shown to be regulated by CLASP, which promotes microtubule polymerization or increases the frequency of microtubule rescues (Bratman and Chang, 2007; Maton et al., 2015; Pereira et al., 2006). Moreover, silencing of the transforming acid coiled-coil (TACC) protein TACC3, which stabilizes microtubules together with XMAP215 (Gergely et al., 2003; Kinoshita et al., 2005; Peset et al., 2005), reduces the rate of spindle elongation by destabilizing midzone microtubules (Lioutas and Vernos, 2013). This strongly suggests that precise regulation of microtubule stability is necessary for proper spindle elongation. Accordingly, in animal cells, the kinesin-4 member Kif4a terminates anaphase B spindle elongation by regulating the inhibition of microtubule polymerization and thus restricting spindle midzone length (Hu et al., 2011). However, how the required velocity of microtubule growth is set precisely and how microtubule dynamics are coordinated with microtubule sliding to allow seamless spindle elongation remains enigmatic.

A straightforward way of coupling microtubule growth and sliding could involve a motor that sets equal sliding and polymerization speeds. Supporting this possibility, *Xenopus* kinesin-5 promotes microtubule polymerization *in vitro* (Chen and Hancock, 2015). The motor has been proposed to induce a curved-to-straight conformational transition of tubulin at microtubule plus-ends, thus promoting microtubule growth by virtue of a stabilizing effect (Chen et al., 2019). This may allow the motor to directly regulate the dynamic behavior of the microtubule tracks which it simultaneously slides. However, the biological significance of the polymerization function of kinesin-5 still has to be investigated *in vivo*.

Like kinesin-5 in most organisms, the kinesin-6 Klp9 promotes anaphase B spindle elongation in fission yeast (Fu et al., 2009; Rincon et al., 2017; Yukawa et al., 2017). Both motors form homotetramers and both slide apart microtubules. Moreover, Klp9 sets the velocity of spindle elongation in a dose-dependent manner (Krüger et al., 2019). Spindles elongate faster as the amount of motors increases at the spindle midzone. Thus, Klp9 alone can set the speed of spindle elongation. We, therefore, wondered if this is solely an effect of the microtubule sliding function, or if Klp9 simultaneously regulates microtubule dynamics.

In the present study, we provide insight into the mechanism of anaphase B spindle elongation, from the regulation of Klp9 localization to the anaphase spindle, to the Klp9-mediated coordination of microtubule sliding and growth. Klp9 recruitment to spindle microtubules is regulated in a dephosphorylation-dependent manner by the known microtubule polymerase Dis1, homologue of XMAP215 and ch-TOG (Matsuo et al., 2016). Further, we provide evidence that, at the spindle, Klp9 performs two functions: the generation of microtubule sliding forces, as shown previously, and the regulation of microtubule dynamics. By using monopolar spindles *in vivo* and *in vitro* reconstitution assays with recombinant Klp9, we could show that the kinesin-6 is a crucial regulator of microtubule growth. In fact, Klp9 increases or decreases the microtubule growth speed depending on the condition. Whereas, at low tubulin concentration, where microtubule growth is comparatively slow, Klp9 increases the microtubule growth speed, at high tubulin concentration, where microtubule growth is fast, Klp9 decreases microtubule growth speed in a dose-dependent manner. This suggests an unconventional mechanism by which Klp9 can promote and block tubulin dimer addition to the microtubule lattice. Eventually, Klp9 is able to set a well-defined microtubule growth velocity. With the dual-function of Klp9 in regulating the microtubule sliding and growth velocity, both processes are inherently coordinated to allow flawless spindle elongation.

## Results

### Monopolar spindles to study microtubule dynamics in anaphase B

We found monopolar spindles to be a useful tool to examine microtubule dynamics not only during early mitotic phases (Costa et al., 2013), but also during anaphase B. Monopolar spindles can be generated by the temperature-sensitive mutant *cut7-24* (Hagan and Yanagida, 1992, 1990). Since the kinesin-5 Cut7 is essential to establish spindle bipolarity during spindle assembly by separating the two spindle poles (Hagan and Yanagida, 1992), inactivation of Cut7 at its restrictive temperature results in the formation of monopolar spindles (Figure 1A).

**Figure 1:**
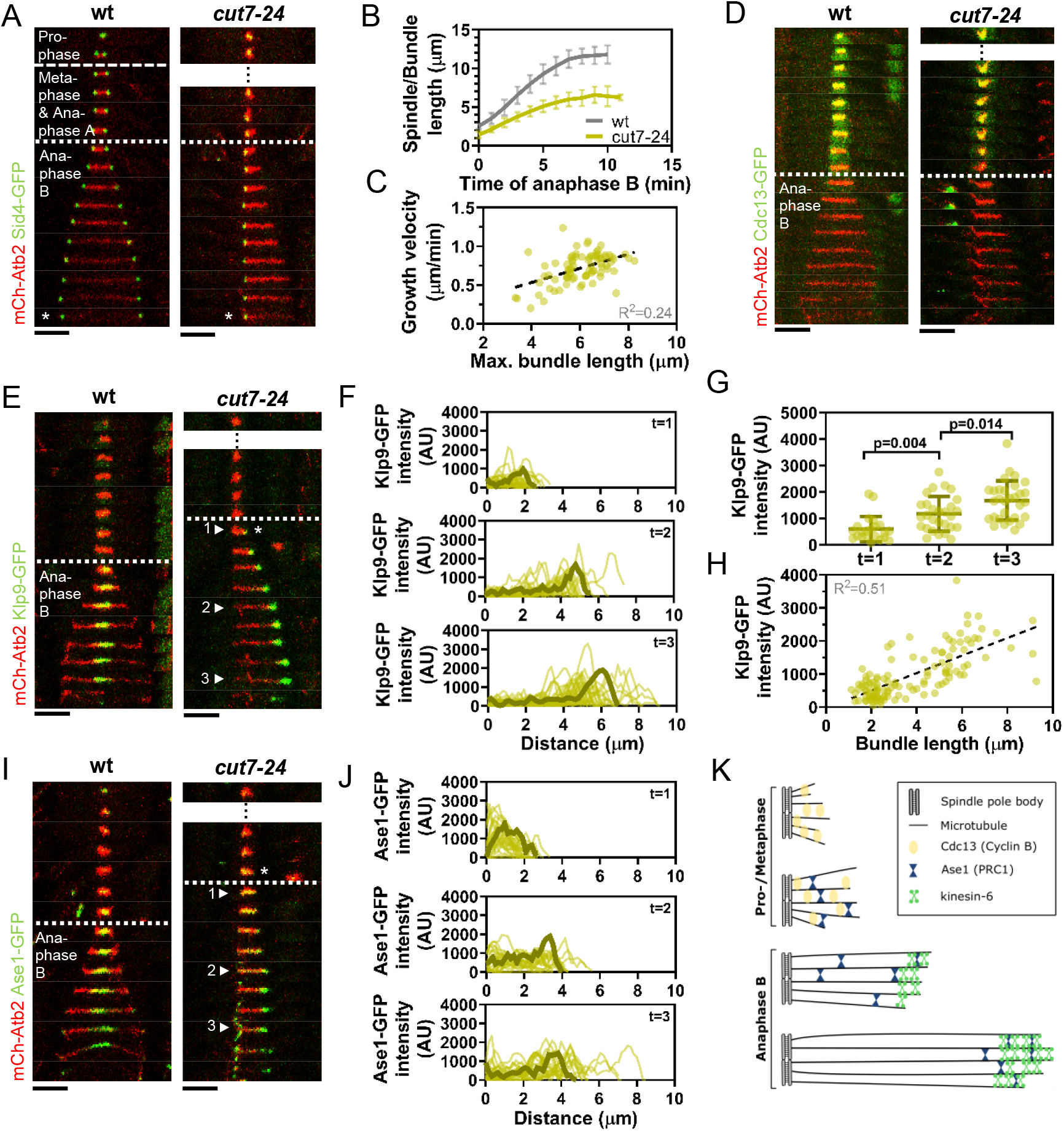
*Cut7-24* monopolar spindle as a tool to study microtubule dynamics during anaphase B. (A) Time-lapse images of wild-type and *cut7-24* cells expressing mCherry-Atb2 (tubulin) and Sid4-GFP (SPBs) at 37°C. The dashed line denotes the transition from prophase to metaphase. Dotted line denotes the transition to anaphase B. Asterisk marks spindle disassembly. Each frame corresponds to 1 min interval. (B) Comparative plot of anaphase B spindle dynamics of wild-type (n=40) or bundle dynamics of *cut-74* cells (n=40) at 37°C. Bold curves correspond to the mean and error bars to the standard deviation. (C) Microtubule bundle growth velocity in *cut7-24* cells plotted against final bundle length. n=79 microtubules (D) Time-lapse images of wild-type and *cut7-24* cells expressing mCherry-Atb2 (tubulin) and Cdc13-GFP (Cyclin B) at 37°C. (E) Time-lapse images of wild-type and *cut7-24* cells expressing mCherry-Atb2 (tubulin) and Klp9-GFP (kinesin-6) at 37°C. Asterisk marks the appearance of the Klp9-GFP signal on the microtubule bundle. Arrowheads 1,2 and 3 correspond to the time points used for linescan analyses at t=1 (first time point of anaphase B), t=2 (5 min after anaphase onset) and t=3 (last time point of anaphase B). (F) Intensity spectra obtained by linescan analysis of Klp9-GFP signals along the microtubule bundles of *cut7-24* cells at time points 1,2 and 3. 0 μm at the x-axis marks the origin of the microtubule bundle at spindle pole bodies. Dark green lines display an exemplary spectrum. (G) Dot plot comparison of the Klp9-GFP intensity on the microtubule bundle of *cut7-24* cells at time points 1 (n=22),2 (n=28) and 3 (n=30). Dark green lines display the mean and standard deviation. P values were calculated using Mann-Whitney U test. (H) Klp9-GFP intensities along microtubule bundles of *cut7-24* cells plotted against microtubule bundle length. n=122 (I) Time-lapse images of wild-type and *cut7-24* cells expressing mCherry-Atb2 (tubulin) and Ase1-GFP at 37°C. Asterisk marks the appearance of the Ase1-GFP signal on the microtubule bundle. Arrowheads 1,2 and 3 correspond to the time points used for linescan analysis at t=1 (first time point of anaphase B), t=2 (5 min after anaphase onset) and t=3 (last time point of anaphase B). Scale bar, 5 μm. (J) Intensity spectra obtained by linescan analysis of Ase1-GFP signals along the microtubule bundles at time points 1,2 and 3. 0 μm at the x-axis marks the origin of the microtubule bundle at spindle pole bodies. Dark green lines display an exemplary spectrum. (K) Model of *cut7-24* monopolar spindles displaying phase I, termed pro-metaphase and phase II corresponding to anaphase B as judged by the absence of the Cdc13-GFP signal and the presence of Klp9-GFP and Ase1-GFP in the long microtubule bundles. In (A),(D), (E) and (I) dotted lines denote the transition to anaphase B. Scale bar, 5 μm. In (C) and (H) data was fitted by linear regression (dashed line), showing the regression coefficient (R^2^) and the slope m. Data from n cells was collected from at least three independent experiments.

Live cell imaging of fission yeast cells expressing α-tubulin (Atb2) linked to mCherry to visualize microtubules and Sid4 linked to GFP to mark spindle pole bodies (SPB) showed the typical three phases of spindle dynamics in wild-type cells: prophase, metaphase and anaphase A, and anaphase B, during which the spindle dramatically elongates until it disassembles (Figure 1A) (Nabeshima et al., 1998). In contrast, the *cut7-24* monopolar spindle underwent only two distinguishable phases (Figure 1A). First, short and rather dynamic microtubules emanated from the two unseparated spindle poles for approximately two hours. Following this phase, one or up to three long microtubule bundles grew from the spindle poles until they disassembled, just before the cell divided (Figure 1A & Figure 1 – figure supplement 1). This extensive growth of microtubule bundles was reminiscent of the microtubule polymerization that is necessary to allow spindle elongation during anaphase B in bipolar spindles. The bundles grew up to an average length of 6.1 ± 1.0 μm, which is slightly longer than half of the average wild-type spindle length (11.6 ± 1.3 μm) (Figure 1B) with an average speed of 0.7 ± 0.2 μm/min, slightly faster than half of the velocity at which the bipolar wild-type spindle elongated (1.2 ± 0.2 μm/min) (Figure 1B). Thus, monopolar spindles appear to behave like half-spindles. Moreover, the growth velocity of individual bundles increased with final bundle length (Figure 1C), a correlation we have previously observed for bipolar anaphase B spindles: longer spindles elongate with respectively higher velocities (Krüger et al., 2019). Together, the second phase of *cut7-24* spindle dynamics resembled anaphase B with the bundle growth velocity matching the speed of spindle elongation in bipolar spindles, and its final length being similar to half the final length of a bipolar anaphase spindle.

To further test if monopolar spindles indeed proceed to anaphase B, we used cyclin B Cdc13 linked to GFP as a marker. Cdc13 disappears from spindle microtubules just before anaphase B onset (Decottignies et al., 2001). Accordingly, we observed fading of the signal on wild-type spindle microtubules and *cut7-24* monopolar spindle microtubules just before the spindle or bundle was elongated (Figure 1D).

Next, we investigated the localization of crucial anaphase B spindle components in monopolar spindles. In bipolar spindles, the sliding motor Klp9 (kinesin-6) localizes to the spindle midzone from anaphase B onset (Fu et al., 2009) (Figure 1E, left panel). In the *cut7-24* monopolar spindle, Klp9-GFP localized to the tip of the microtubule bundle once it started to elongate (Figure 1E, right panel). The Klp9-GFP intensity profile obtained at three different time points revealed that Klp9-GFP preferentially localized to the tip of the bundle from the first time point of anaphase B (Figure 1F) and the intensity increased with anaphase B progression (Figure 1F & 1G). Hence, while the bundle was growing Klp9 accumulated at its tip. In general, we observed a strong correlation of the Klp9-GFP intensity at the bundle tip and the bundle length (Figure 1H).

Last, we analyzed the localization of the conserved microtubule bundler Ase1. Ase1 crosslinks the antiparallel overlapping microtubules at the spindle center and stabilizes the spindle structure (Janson et al., 2007; Loiodice et al., 2005; Yamashita et al., 2005). In the wild-type spindle, Ase1-GFP localized to the spindle midzone just before anaphase onset and onward, as previously reported (Loiodice et al., 2005) (Figure 1I, left panel). In the monopolar spindle, Ase1-GFP similarly localized to the spindle just before the microtubule bundle started to grow (Figure 1I, right panel). The Ase1-GFP intensity profiles showed that the signal was spread all along the bundle at early time points of bundle elongation (Figure 1I & J; timepoint 1), and accumulated at the bundle tip at later time points (Figure 1I & J; timepoint 2 & 3).

Taken together, besides unseparated spindle poles, *cut7-24* monopolar spindles proceed to anaphase B, during which long microtubule bundles are polymerized, as judged by the absence of the Cdc13-GFP signal, and the presence of Klp9-GFP and Ase1-GFP on the microtubule bundles (Figure 1K). Moreover, the bundle growth velocity equals half of the spindle elongation velocity of bipolar spindles, suggesting that microtubule dynamics are not altered in the mutant. Therefore, monopolar spindles constitute a suitable tool to study microtubule dynamics during anaphase B.

### The kinesin-6 Klp9 affects microtubule growth during anaphase B

Using monopolar spindles, we could now analyze the effect of Klp9 on microtubule dynamics. As reported earlier (Fu et al., 2009), deletion of *klp9* strongly decreased the speed of bipolar spindle elongation in anaphase B (Figure 2A). Deletion of *klp9* in the *cut7-24* background prevented the formation of long microtubule bundles (Figure 2B). While anaphase B microtubule bundles reached a maximum length of 6.1 ± 1.0 μm in presence of Klp9, the bundles only grew up to 2.7 ± 0.5 μm upon *klp9* deletion (Figure 2C). Moreover, the growth velocity of the bundles was strongly reduced upon *klp9* deletion *(cut7-24:* 0.7 ± 0.2 μm/min; *cut7-24 klp9Δ:* 0.1 ± 0.1 μm/min) (Figure 2C). To test if this effect is dose-dependent, we used a shut-off strain, in which the expression of *klp9* is strongly reduced, but some Klp9 molecules are still present. Indeed, the phenotype was slightly less dramatic as compared to the deletion of *klp9* (Figure 1C). The bundles reached a maximum length of 3.2 ± 0.8 μm, and grew with an average velocity of 0.2 ± 0.2 μm/min. This suggests that Klp9 is involved in the regulation of microtubule growth during anaphase B.

**Figure 2:**
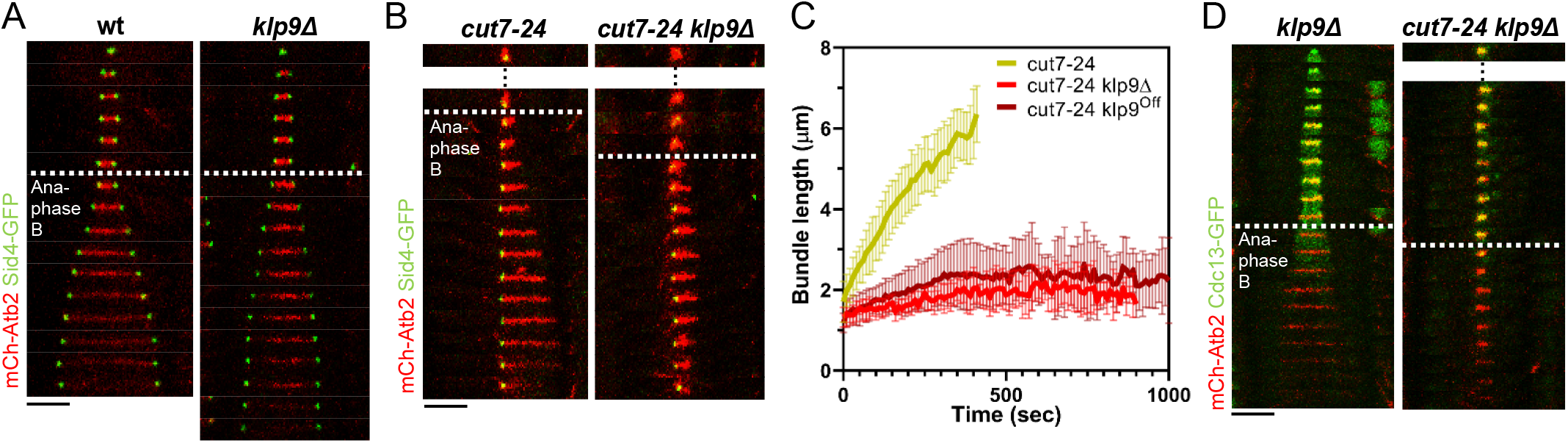
Klp9 promotes microtubule growth during anaphase B in monopolar spindles. (A) Time-lapse images of wild-type and *klp9Δ* cells expressing mCherry-Atb2 (tubulin) and Sid4-GFP (SPBs) at 37°C. (B) Time-lapse images of *cut7-24* and *cut7-24 klp9Δ* cells expressing mCherry-Atb2 (tubulin) and Sid4-GFP (SPBs) at 37°C. (C) Comparative plot of microtubule bundle dynamics in *cut7-24* (n=40), *cut7-24 klp9Δ* (n=40) and *cut7-24 klp9^Off^* (n=40) at 37°C. Bold curves correspond to the mean and error bars to the standard deviation. (D) Time-lapse images of *klp9Δ* and *cut7-24 klp9Δ* cells expressing mCherry-Atb2 (tubulin) and Cdc13-GFP (Cyclin B) at 37°C. In (A-B) and (D) each frame corresponds to 1 min interval. Dotted lines denote the transition to anaphase B. Scale bar, 5 μm. Data from n cells was collected from at least three independent experiments.

However, Klp9 has been previously implicated in the regulation of the metaphase-to-anaphase transition (Meadows et al., 2017). We thus had to rule out that the observed reduction of microtubule growth is not a consequence of an impaired anaphase B transition in monopolar spindles upon *klp9* deletion. To do so, we analyzed the localization of Cdc13-GFP. In a bipolar *klp9Δ* spindle, Cdc13-GFP disappeared from the spindle just before it started to elongate (Figure 2D). In the *cut7-24 klp9Δ* mutant, Cdc13-GFP also disappeared from the spindle microtubules, but still no long microtubule bundle was formed (Figure 2D). Hence, the observed phenotype can be attributed to the Klp9 function in microtubule growth regulation during anaphase B.

### The XMAP215 family member Dis1 displays a similar effect on microtubule bundle growth as Klp9

We hypothesized, that Klp9 could either regulate microtubule dynamics by itself or indirectly through another microtubule-associated protein (MAP). To probe the involvement of other proteins, we analyzed the impact of the deletion of several candidates on microtubule bundle growth. Namely, the conserved microtubule bundler Ase1, which has been proposed to recruit Klp9 and other spindle components, such as CLASP to the mitotic spindle (Bratman and Chang, 2007; Fu et al., 2009); the EB1 homolog Mal3, which impacts microtubule dynamics *in vitro* (Bieling et al., 2007; Des Georges et al., 2008; Katsuki et al., 2009; Matsuo et al., 2016); the two members of the XMAP215 family, Dis1 and Alp14, which act as microtubule polymerases (Al-Bassam et al., 2012; Matsuo et al., 2016); and the TACC protein Alp7, which interacts with Alp14 and enhances its polymerase activity (Hussmann et al., 2016; Sato et al., 2004).

The deletion of *ase1, mal3, alp14*, and *alp7* still allowed the formation of comparatively long microtubule bundles in the *cut7-24* background (Figure 3A & B). In all cases, the growth velocity of microtubule bundles was significantly reduced (Figure 3B & C). However, only the deletion of *dis1* led to a decrease of the bundle growth velocity comparable to the deletion or shut-off condition of *klp9* (Figure 3B & C). Moreover, maximum bundle length was reduced upon deletion of *ase1, dis1*, and *alp7*, with the strongest decrease observed upon *dis1* deletion (Figure 3D).

**Figure 3:**
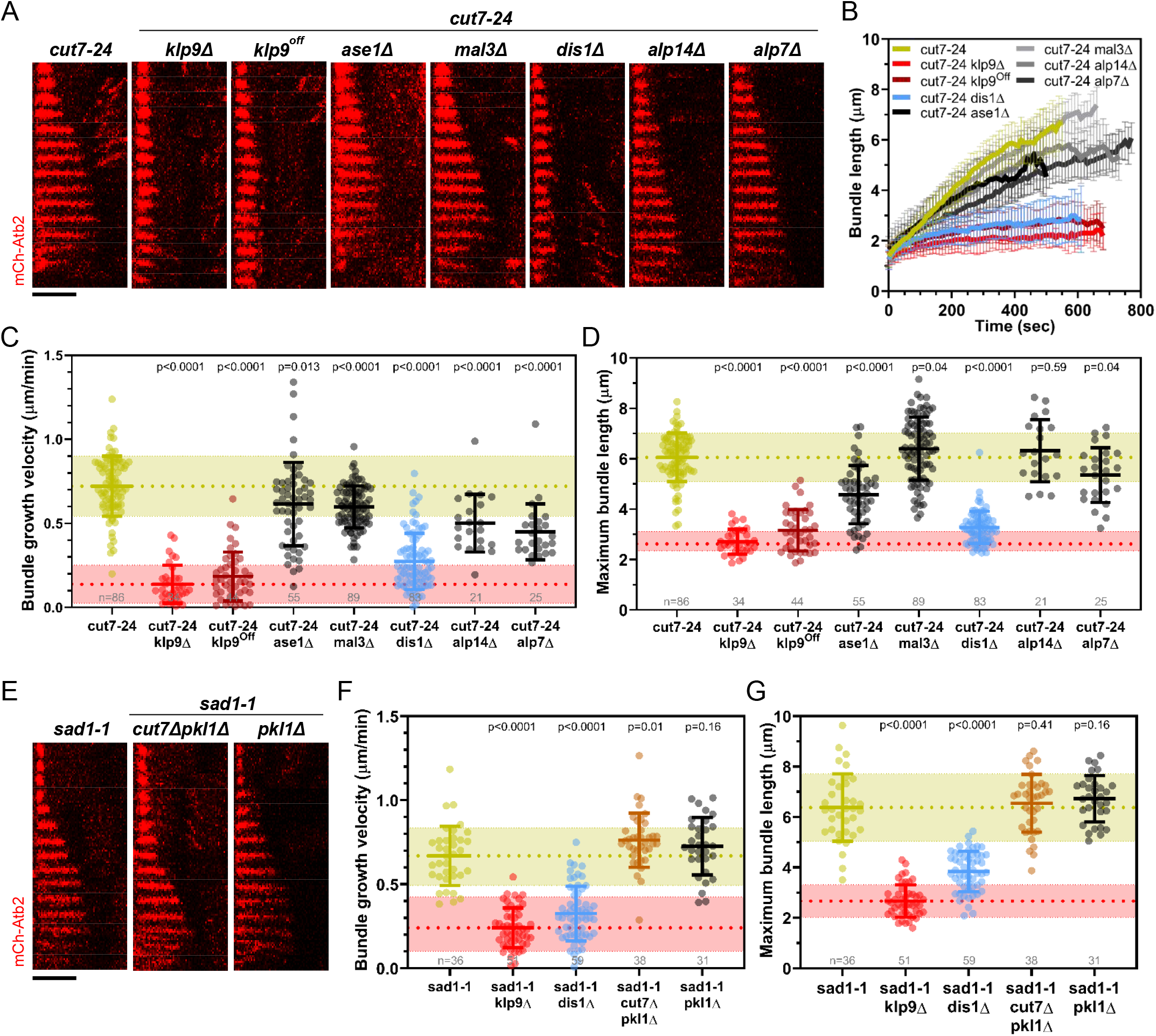
Deletion of *dis1* decreases microtubule bundle growth velocity and bundle length during anaphase B in monopolar spindles. (A) Time-lapse images of *cut7-*24, *cut7-24 klp9Δ, cut7-24 klp9^Off^, klp9^Off^ ase1Δ, cut7-24 mal3Δ, cut7-24 dis1Δ, cut7-24 alp14Δ* and *cut7-24 alp7Δ* cells expressing mCherry-Atb2 (tubulin) at 37°C. (B) Comparative plot of microtubule bundle dynamics in *cut7-24* (n=40), *cut7-24 klp9Δ* (n=40), *cut7-24 klp9^Off^*(n=40), *cut7-24 ase1Δ* (n=39), *cut7-24 mal3Δ* (n=40), *cut7-24 dis1Δ* (n=39), *cut7-24 alp14Δ* (n=20) and *cut7-24 alp7Δ* cells (n=25) at 37°C. Bold curves correspond to the mean and error bars to the standard deviation. (C) Dot plot comparison of microtubule bundle growth velocity during anaphase B. (D) Dot plot comparison of maximum microtubule bundle length during anaphase B. (E) Time-lapse images of *sad1-1, sad1-1 cut7Δpkl1Δ* and *sad1-1 pkl1Δ* cells expressing mCherry-Atb2 (tubulin) at 37°C. (F) Dot plot comparison of microtubule bundle growth velocity during anaphase B. (G) Dot plot comparison of maximum microtubule bundle length during anaphase B. In (A) and (E) Each frame corresponds to 1 min interval. Scale bar, 5 μm. In (C-D) and (F-G) lines correspond to mean and standard deviation. P values were calculated using Mann-Whitney U test. Data from n cells was collected from at least three independent experiments.

We note that the deletion of *alp14* and *alp7* often resulted in the restoration of spindle bipolarity. This is in agreement with the model, that a balance between microtubule dynamics and microtubule crosslinking is crucial for the establishment of spindle bipolarity (Yukawa et al., 2019b). Here, we analyzed the fraction of cells in which spindle bipolarity could not be restored, and spindles remained monopolar.

Taken together, of the tested candidates, the deletion of *dis1* gave rise to a similar phenotype as the deletion or shut-off condition of *klp9*. Similar to the deletion of *klp9*, upon *dis1* deletion the transition to anaphase B is not impaired, as indicated by the disappearance of the Cdc13-GFP signal (Figure 3 – figure supplement 1). Thus, like Klp9, Dis1 seems to be involved in the regulation of microtubule growth during anaphase B and may act in the same pathway.

Besides, we examined a possible involvement of the kinesin-5 Cut7 and CLASP Cls1 (also called Peg1). Cut7 has been shown to promote spindle elongation like Klp9, even though to a lower extent (Rincon et al., 2017; Yukawa et al., 2019b), and a dimeric construct of the *X. laevis* kinesin-5 Eg5 promotes microtubule growth *in vitro* (Chen et al., 2019; Chen and Hancock, 2015). To test the role of Cut7 we used the *sad1-1* temperature-sensitive mutant to generate monopolar spindles (Hagan and Yanagida, 1995). *Cut7* deletion was performed in the background of *pkl1* (kinesin-14) deletion, since the deletion of the kinesin-5 alone is lethal (Olmsted et al., 2014; Rincon et al., 2017; Syrovatkina and Tran, 2015; Yukawa et al., 2018). *Sad1-1* monopolar spindles also assembled long microtubule bundles during anaphase B (Figure 3E, Figure 3 – figure supplement 2). Unlike the deletion of *klp9* or *dis1*, the deletion of *cut7 (cut7Δpkl1Δ)* led to a slight increase of the growth velocity (Figure 3F) and no significant change of maximum bundle length (Figure 3G). The modest acceleration of bundle growth seemed to be a result of the absence of Cut7 and not Pkl1, since the deletion of *pkl1* alone did not affect growth velocity significantly (Figure 3F). Furthermore, we investigated the localization of Cut7-GFP in the *sad1-1* mutant. While, Klp9-GFP was detected at the tip of microtubule bundles, Cut7-GFP localized only to the unseparated spindle poles (Figure 3 – figure supplement 2). Thus, even though both motors have been reported to be plus-end directed bipolar kinesins, and both localize to spindle microtubules in a bipolar spindle (Fu et al., 2009; Hagan and Yanagida, 1992), only Klp9 localizes to the tip of the microtubule bundles in monopolar spindles, and promotes their growth during anaphase B.

Last, we examined if the phenotype upon *klp9* deletion could stem from an interaction with the fission yeast CLASP Cls1. Cls1 localizes to the spindle midzone during anaphase B and promotes microtubule rescues, thus preventing microtubules from depolymerizing up to the spindle poles and preserving spindle stability (Bratman and Chang, 2007). Since deletion of *cls1* is lethal, we tested a potential interaction with Klp9 during anaphase B by expressing Cls1-3xGFP in wild-type and *klp9* deleted cells. In bipolar spindles, Cls1-3xGFP localized to the spindle midzone in presence or absence of Klp9 (Figure 3 – figure supplement 3) with similar intensities throughout anaphase B (Figure 3 – figure supplement 4). Moreover, in monopolar *cut7-24* spindles, Cls1-3xGFP, unlike Klp9-GFP, did not localize to the bundle tip but was rather spread all along the bundle (Figure 3 – figure supplement 5). Therefore, we conclude that Cls1 is not involved in the underlying mechanism of Klp9-mediated microtubule growth during anaphase B.

### Dis1 regulates the recruitment of Klp9 to the anaphase spindle

Of the tested candidates, only the XMAP215 family member Dis1 emerged as a possible candidate that acts in the pathway with Klp9. To probe this, we examined its role during anaphase B spindle elongation in bipolar spindles. Similar to the deletion of *klp9*, the deletion of *dis1* decreased the spindle elongation velocity (Figure 4A & B). Simultaneous deletion of both proteins did not display an additive effect, and decreased spindle elongation velocity to not significantly different values as the deletion of individual proteins (Figure 4A & B), suggesting that Klp9 and Dis1 act in the same pathway. Furthermore, final spindle length is reduced in all three mutants (Figure 4C), further indicating an effect of Dis1 and Klp9 on microtubule growth in bipolar spindles. Dis1 was previously shown to be a microtubule polymerase (Matsuo et al., 2016). It was thus tempting to think that Klp9 transports Dis1 to the plus-ends of microtubules. However, Dis1-EGFP localization to spindle poles and the lateral spindle microtubule lattice, and Dis1-EGFP intensity on the spindle was not altered in absence of Klp9 (Figure 4 – figure supplement 1). Moreover, in monopolar spindles, Dis1-EGFP localized to the unseparated spindle poles and disperse along the microtubule bundle (Figure 4 – figure supplement 2), but did not accumulate at the bundle. These results suggest that Klp9 does neither regulate the recruitment of Dis1 nor its localization along the anaphase spindle in bipolar or monopolar spindles.

**Figure 4:**
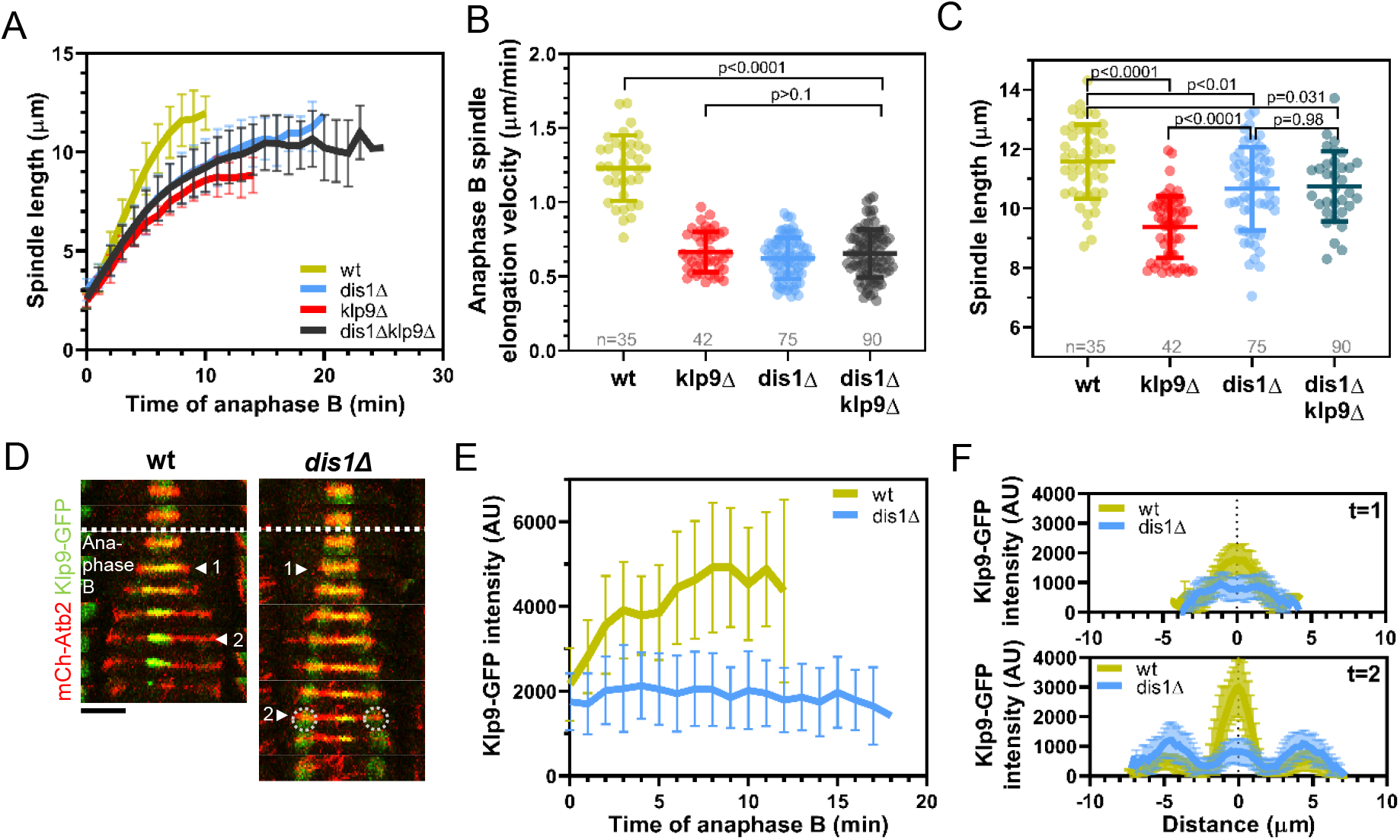
*Dis1* deletion impairs Klp9 recruitment to the anaphase spindle. (A) Comparative plot of anaphase B spindle dynamics of wild-type (n=35), *dis1Δ* (n=75), *klp9Δ* (n=38) and *dis1Δklp9Δ* (n=90) at 25°C. (B) Dot plot comparison of spindle elongation velocity in wild-type, *dis1Δ, klp9Δ* and *dis1Δklp9Δ* cells. (C) Dot plot comparison of final anaphase B spindle length in wild-type, *dis1Δ, klp9Δ* and *dis1Δklp9Δ* cells. (D) Time-lapse images of wild-type and *dis1Δ* cells expressing mCherry-Atb2 (tubulin) and Klp9-GFP at 25°C. Arrowheads 1 and 2 correspond to the time points used for linescan analysis at t=1 (2 min after anaphase B onset), t=2 (2 min before spindle disassembly). Circles mark the Klp9 pool that remained in the nucleoplasm. Scale bar, 5 μm. (E) Comparative plot of Klp9-GFP intensity throughout anaphase B spindle elongation of wild-type (n=30) and *dis1Δ* cells (n=33). (F) Intensity spectra obtained by linescan analysis of Klp9-GFP signals along the anaphase spindle at early (t=1) and late anaphase (t=2). In (A), (E) and (F) bold curves correspond to the mean and error bars to the standard deviation. In (B) and (C) lines correspond to mean and standard deviation. Data from n cells was collected from at least three independent experiments.

Therefore, we wondered if Dis1 could recruit Klp9. Indeed, we observed that the Klp9-GFP signal at the spindle midzone was strongly diminished upon *dis1* deletion (Figure 4D). While in presence of Dis1 the intensity of Klp9-GFP increased with progressing spindle elongation, until a plateau was reached in late anaphase, the intensity remained low in *dis1Δ* cells (Figure 4E). This is not a consequence of overall lower Klp9 levels, since the total intensity of Klp9-GFP (measured at mitosis onset in the nucleoplasm, where Klp9 is localized before anaphase onset) was not significantly altered (Figure 4 – Figure supplement 3). Intensity profiles along the spindle at early and late anaphase B showed that the midzone localization of Klp9-GFP was not impaired in absence of Dis1 (Figure 4F). The increased intensity of Klp9-GFP close to the spindle poles in late anaphase in *dis1Δ* (Figure 4F, t=2) corresponded to the pool of Klp9-GFP that remained in the nucleoplasm and was thus not recruited to the spindle (Figure 4D, circles). To further exclude the possibility, that the decreased Klp9-GFP intensity could be a result of a decreased spindle microtubule number, and thus fewer binding-sites for Klp9 upon *dis1* deletion, we normalized the Klp9-GFP intensity with the mCherry-Atb2 intensity. This relative Klp9 concentration at the midzone increased in wild-type cells, according to the increasing Klp9-GFP intensity, but remained low in *dis1Δ* cells (Figure 4 - figure supplement 4). Hence, the impaired localization of Klp9-GFP did not stem from differences in microtubule number. Together, the results indicate that Dis1 regulates the recruitment of the motor to the anaphase spindle.

To further probe this idea, we slightly overexpressed *dis1* by inserting the thiamine repressible nmt promoter *pnmt81* upstream of the *dis1* open reading frame. The intensity of Klp9-GFP was increased upon overexpression of *dis1* throughout anaphase B at the midzone while the total Klp9-GFP intensity was not significantly different (Figure 4 – figure supplement 5). Thus, the microtubule polymerase Dis1 regulates the recruitment of the kinesin-6 motor Klp9 to the anaphase B spindle in a dose-dependent manner.

### Dis1 regulates the recruitment of Klp9 in a dephosphorylation-dependent manner

Dis1 localization is regulated in a phosphorylation-dependent manner during mitosis. Phosphorylation of Dis1 by Cdc2, homolog of Cdk1, promotes its localization to kinetochores during metaphase (Aoki et al., 2006), where it is involved in sister chromatid separation (Nabeshima et al., 1995; Ohkura et al., 1988). Dephosphorylation of the Cdc2-phosphosites at the metaphase-to-anaphase transition results in disappearance of Dis1 from kinetochores and its relocalization to the lateral microtubule lattice of anaphase spindles (Aoki et al., 2006). At this location Dis1 may be required for bundling parallel microtubules (Roque et al., 2010). The phosphatase that mediates Dis1 dephosphorylation is not known. Besides, Klp9 has also been shown to be phosphorylated by Cdc2 at mitosis onset (Fu et al., 2009). Dephosphorylation at the metaphase-anaphase transition by Clp1, a homolog of Cdc14, is suggested to trigger Klp9 recruitment to the anaphase spindle (Fu et al., 2009). We thus wondered if Dis1 is also dephosphorylated by Clp1, and further, if this is required for Dis1-mediated Klp9 recruitment.

To test this idea, we used a strain expressing Klp9-mCherry and Dis1-GFP in presence or absence of Clp1. Upon *clp1* deletion Dis1-EGFP could only be detected at spindle poles, and not on spindle microtubules upon anaphase B onset (Figure 5A - C), resulting in a decreased Dis1-EGFP intensity along the spindle throughout anaphase B (Figure 5D). This localization pattern of Dis1-EGFP was equal to a previously analyzed phosphomimetic Dis1 mutant (Aoki et al., 2006), indicating that in absence of Clp1, Dis1 remained phosphorylated. Moreover, *clp1* deletion resulted in a significant reduction of the Klp9-mCherry intensity at the spindle midzone (Figure 5A, C & E). Consequently, the spindle elongation velocity in anaphase B was reduced upon *clp1* deletion (Figure 5 – figure supplement 1). Thus, Klp9 and Dis1 are dephosphorylated at the metaphase-anaphase transition by Clp1, allowing Dis1 to bind to the parallel spindle microtubule lattice, which may subsequently promote Klp9 recruitment to antiparallel midzone microtubules (Figure 5F).

**Figure 5:**
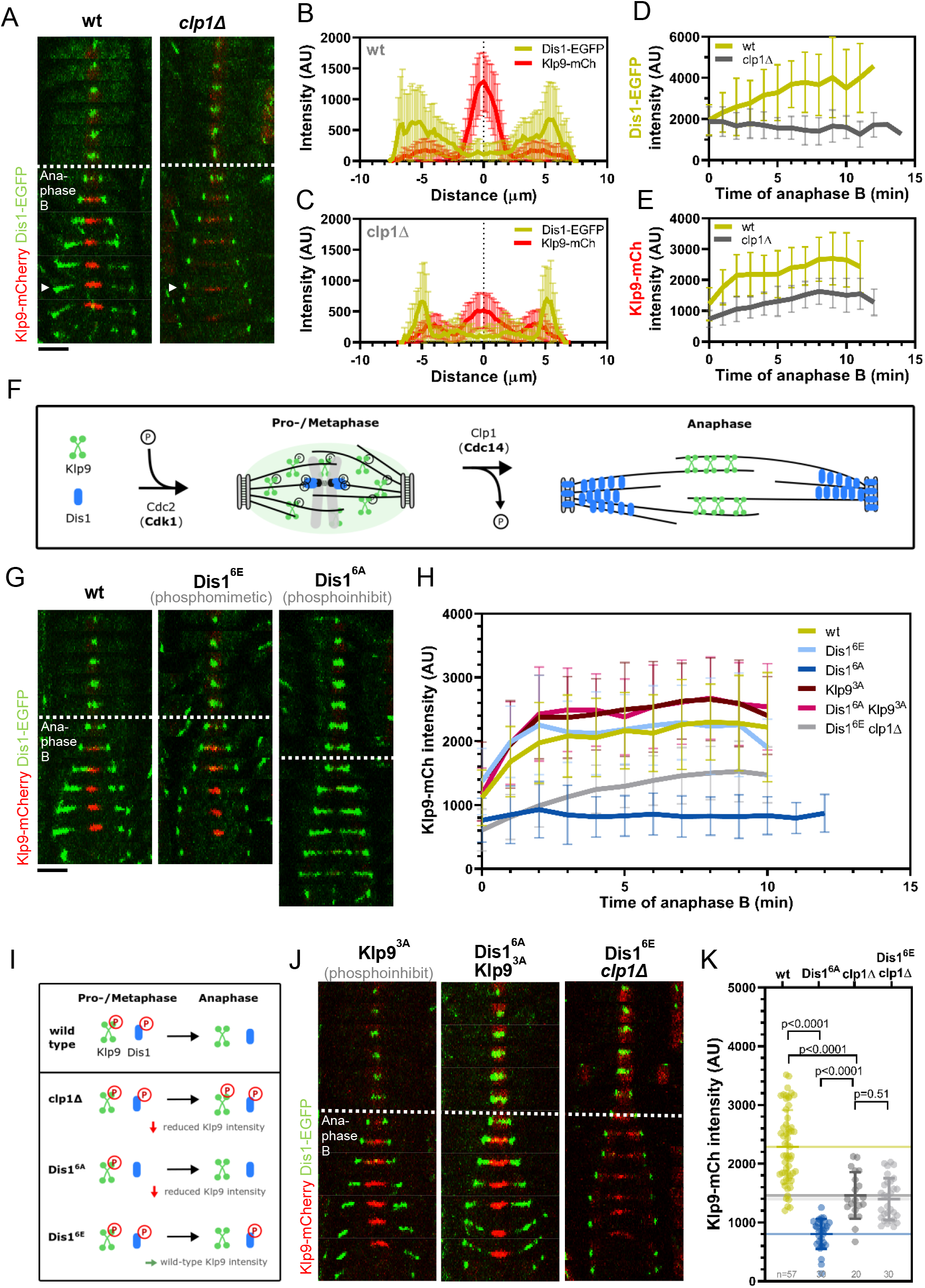
Phospho-dependent regulation of Dis1-mediated Klp9 recruitment. (A) Time-lapse images of wild-type and *clp1Δ* cells expressing Klp9-mCherry and Dis1-EGFP at 25°C. Arrowhead depicts the time point used for linescan analysis (2 min before spindle disassembly). (B) Intensity spectra obtained by linescan analysis of Dis1-EGFP and Klp9-mCherry signals along the anaphase spindle at late anaphase in wild-type cells (n=30). X = 0 μm equals the cell center. (C) Intensity spectra obtained by linescan analysis of Dis1-EGFP and Klp9-mCherry signals along the anaphase spindle at late anaphase in *clp1Δ* cells (n=30). (D) Comparative plot of Dis1-EGFP intensity throughout anaphase B spindle elongation of wild-type (n=30) and *clp1Δ* (n=30). (E) Comparative plot of Klp9-mCherry intensity throughout anaphase B spindle elongation of wild-type (n=30) and *clp1Δ* (n=30). (F) Model of phosphorylation-dependent localization of Klp9 (green) and Dis1 (blue) throughout mitosis mediated by the cyclin-dependent kinase Cdc2 (homolog of Cdk1) and the phosphatase Clp1 (homolog of Cdc14). (G) Time-lapse images of cells expressing wild-type Dis1-GFP, phosphomimetic Dis1^6E^-GFP or phosphoinhibit Dis1^6A^-GFP together with Klp9-mCherry at 25°C. (H) Comparative plot of Klp9-mCherry intensity throughout anaphase B spindle elongation of wild-type cells (n=30) and cells expressing Dis1^6E^-GFP (n=30) Dis1^6A^-GFP (n=30), Klp9^3A^-mCherry (n=30), Dis1^6A^-EGFP and Klp9^SA^-mCherry (n=30) and Dis1^6E^ *clp1Δ* cells (n=30). (I) Summary of the results obtained upon *clp1* deletion and expression of phosphoinhibit Dis1^6A^ or phosphoimimetic Dis1^6E^. (J) Time-lapse images of cells phosphoinhibit Klp9^3A^-mCherry with wild-type Dis1-GFP, phosphoinhibit Klp9^3A^-mCherry with phosphoinhibit Dis1^6A^-EGFP, or Dis1^6A^-EGFP in *clp1Δ* cells at 25°C. (K) Dot plot comparison of the Klp9-mCherry intensity (AU) in wild-type, Dis1^6A^, *clp1Δ* and Dis1^6E^ *clp1Δ* cells. Lines correspond to mean and standard deviation. Long lines depict the mean for each cell type. P values were calculated using Mann-Whitney U test. In (A), (G) and (J) Each frame corresponds to 2 min interval. Dotted lines denote the transition to anaphase B. Scale bar, 5 μm. In (B-E) and (H) bold curves correspond to the mean and error bars to the standard deviation. Data from n cells was collected from at least three independent experiments.

However, Clp1 also dephosphorylates the Ase1, and dephosphorylation of Ase1 and Klp9 has been proposed to allow their physical interaction, and eventually the recruitment of Klp9 to the spindle by Ase1 (Fu et al., 2009). Hence, we examined if the observed reduction of the Klp9-mCherry intensity in *clp1Δ* did not stem from prevented Klp9-Ase1 interaction. Focusing on intact spindles, we could not detect mis-localisation of Klp9 or Dis1, nor a reduction of the Klp9-mCherry intensity upon *ase1* deletion (Figure 5 – figure supplement 2 & 3). Thus, Ase1 does not appear to recruit Klp9 to the spindle.

To determine if the decreased recruitment of Klp9 upon *clp1* deletion was a result of the prevented Dis1 dephosphorylation, we took advantage of previously generated phosphomimetic and phosphoinhibit mutants of Dis1. The six consensus Cdc2-phosphosites of Dis1 (279T, 293S, 300S, 551S, 556S, and 590S) were either changed to alanine, to create a phosphoinhibit mutant (Dis1^6A^), or to glutamate, to obtain a phosphomimetic version of Dis1 (Dis1^6E^) (Aoki et al., 2006). If dephosphorylation of Dis1 would be necessary for Klp9 recruitment, we expected the expression of Dis1^6E^-GFP to result in reduced Klp9-mCherry intensities, and expression of Dis1^6A^-GFP to lead to increased Klp9-mCherry intensities at the spindle midzone. Contrarily, we found that the intensity of Klp9-mCherry was not altered in cells expressing the phosphomimetic Dis1^6E^-GFP, but was strongly reduced in cells expressing the phosphoinhibit Dis1^6A^-GFP compared to the expression of wild-type Dis1-GFP (Figure 5G & H). This indicates that the recruitment of Klp9 by Dis1 is independent of the localization of Dis1 to the spindle microtubule lattice but it depends on its phosphorylation: For proper Klp9 recruitment, Dis1 has to be phosphorylated at Cdc2-phosphosites. Taken together, *clp1* deletion resulted in the presence of phosphorylated Klp9 and Dis1 in anaphase B, and reduced recruitment of Klp9 (Figure 5I). This impaired Klp9 recruitment appeared not to stem from the presence of phosphorylated Dis1 during anaphase B, since the expression of the phosphomimetic Dis1^6E^-GFP did not result in decreased Klp9-mCherry intensity at the spindle midzone. Thus, it may originate from abolishing the dephosphorylation of Klp9 by clp1, as previously suggested (Fu et al., 2009). Moreover, the expression of phosphoinhibit Dis1^6A^-GFP strongly reduced Klp9 recruitment to the anaphase spindle. This indicates, that the phosphorylated form of Dis1 regulates the dephosphorylation of Klp9, which eventually triggers proper Klp9 recruitment to the anaphase spindle (Figure 5I). Since Dis1 is phosphorylated only in pro- and metaphase but not in anaphase B when Klp9 recruitment occurs, it could thus be that phosphorylated Dis1 initiates a pathway just before transition to anaphase B that eventually leads to Klp9 dephosphorylation at anaphase onset.

To probe this hypothesis, we decided to test if the expression of a phosphoinhibit version of Klp9 could rescue the decreased Klp9 recruitment observed upon expression of phosphoinhibit Dis1^6A^-GFP. Previously, three Cdc2-dependent phosphorylation sites of Klp9 (S598, S605, and S611) have been mutated to alanine to obtain phosphoinhibit Klp9 (Klp9^3A^) (Fu et al., 2009). Indeed, the intensity of Klp9^3A^-mCherry in cells expressing Dis1^6A^-GFP was not significantly different as compared to cells expressing wild-type Dis1-GFP and Klp9^3A^-mCherry. Thus, the reduced Klp9 recruitment upon expression of Dis1^6A^-GFP could be rescued by simultaneous expression of Klp9^3A^-mCherry. Hence, Dis1 regulates the localization of Klp9 to anaphase spindles via Klp9 dephosphorylation.

Subsequently, the question arose if Dis1 regulates Klp9 dephosphorylation through Clp1. First, interaction of phosphorylated Dis1 with Clp1 could activate Clp1 and promote Dis1 dephosphorylation. Second, activated Clp1 may promote Klp9 dephosphorylation. In this case, the expression of phosphomimetic Dis1^6E^-GFP in cells deleted for *clp1* would result in a similar decrease of the Klp9-mCherry intensity as compared to the expression of phosphoinihbit Dis1^6A^-GFP. The Klp9-mCherry intensity was decreased upon deletion of *clp1* in cells expressing phosphomimetic Dis1^6E^-GFP (Figure 5H & J). However, as in *clp1Δ* cells, in Dis1^6E^-GFP *clp1Δ* cells, the Klp9-mCherry intensity increased throughout anaphase B, unlike in cells expressing Dis1^6A^-GFP, where the intensity of Klp9-mCherry remained constant (Figure 5E & H). As a result, Klp9-mCherry intensity became significantly higher in Dis1^6E^-GFP *clp1Δ* and *clp1Δ* cells, with respect to Dis1^6A^-GFP cells (Figure 5H & K). We noticed that the sum of the Klp9-mCherry intensity in cells expressing Dis1^6A^-GFP (804 ± 255 AU) and Dis1^6E^-GFP *clp1Δ* (1398 ± 355 AU) or *clp1Δ* cells (1460 ± 396 AU) yields a similar value as the intensity in wild-type cells (2287 ± 629 AU). Hence, the dephosphorylation of Klp9, which allows binding to spindle microtubules, appears to be regulated by two pathways: one relying on phosphorylated Dis1 (~65%) and one on Clp1 (~35%).

### Klp9 sets the microtubule growth velocity *in vitro*

Since Dis1 appeared to act upstream of Klp9, the effect of Klp9 on microtubule polymerization in monopolar spindles may arise from the motor function itself. To probe this hypothesis, we examined the effect of recombinant full-length Klp9 (Figure 6 – figure supplement 1) on dynamic microtubules *in vitro*. The functionality of the kinesin-6 was tested using a microtubule gliding assay (Figure 6A). Klp9 was immobilized on a glass coverslip via a His6-antibody and red ATTO-647-labeled taxol-stabilized microtubules polymerized from porcine tubulin were added to the flowchamber (Figure 6A). Using Total Internal Reflection Fluorescence (TIRF)-microscopy, microtubule motion was observed. Microtubules moved with an average velocity of 2.4 ± 0.4 μm/min (Figure 6B & C) which was consistent with previously observed Klp9-mediated microtubule gliding velocities (Yukawa et al., 2019a).

**Figure 6:**
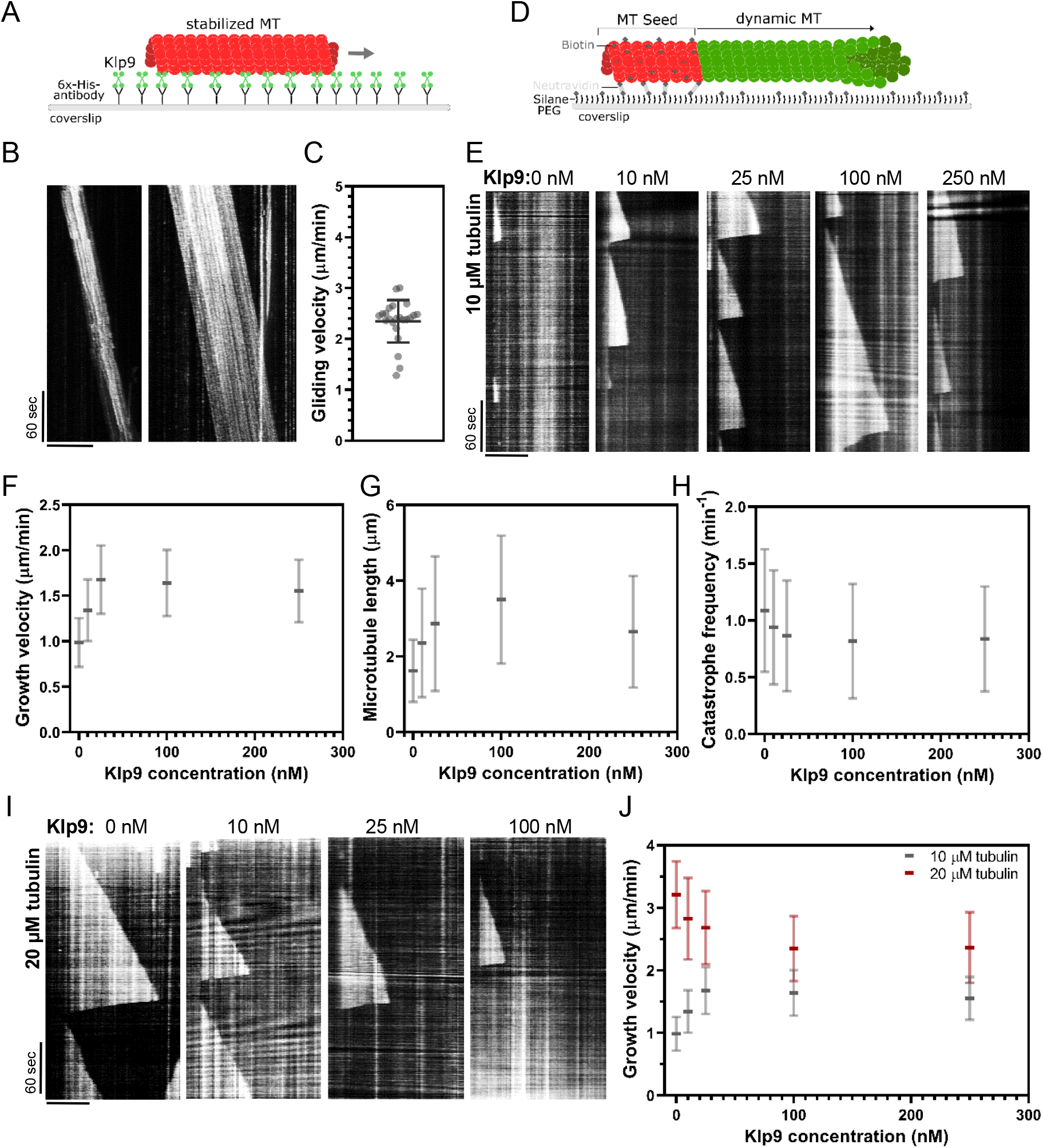
Recombinant Klp9 regulates the microtubule growth velocity *in vitro*. (A) Schematic set up of the microtubule gliding assay. His6 antibodies are shown in gray, Klp9 molecules in green and taxol-stabilized microtubules in red. (B) Kymographs of gliding microtubules with the time on the y-axis and space on the x-axis. Scale bar, 10 μm. (C) Dot plot of microtubule gliding velocities. (D) Schematic of microtubule polymerization assay: GMPCPP-stabilized, ATTO-647 labeled microtubule seeds are shown in red and the dynamic microtubule grown from free tubulin (80% unlabeled, 20% ATTO-488-labeled tubulin) in green. (E) Kymographs of dynamic microtubules grown in presence of 10 μM free tubulin and 0, 10, 25, 100 and 250 nM Klp9. Scale bar, 5 μm. (F) Microtubule growth velocity (μm/min) shown as a function of the Klp9 concentration (nM); n=190-325 microtubules per condition. (G) Microtubule length (μm) shown as a function of the Klp9 concentration (nM); n=157-325 microtubules per condition. (H) Catastrophe frequency (min^-1^) shown as a function of the Klp9 concentration (nM); n=65-112 microtubules per condition. (I) Kymographs of dynamic microtubules grown in presence of 20 μM free tubulin (80% unlabeled, 20% ATTO-488-labeled tubulin) and 0, 10, 25 and 100 nM Klp9. Scale bar, 5 μm. (J) Microtubule growth velocity (μm/min) measured in presence of 10 μM or 20 μM free tubulin shown as a function of the Klp9 concentration (nM); n=190-399 microtubules per condition. For (F-H) and (J) mean values and standard deviations are shown. Data from n microtubules was collected from at least three independent experiments.

Subsequently, we examined the effect of Klp9 on dynamic microtubules growing from biotin- and ATTO-647-labeled, GMPCPP-stabilized microtubule seeds (Figure 6D). In presence of 10 μM free tubulin (80% unlabeled, 20% ATTO-488 labeled), addition of Klp9 increased the growth velocity in a dose-dependent manner from 1.0 ± 0.3 μm/min, in absence of Klp9, to 1.7 ± 0.4 μm/min, in presence of 25 nM Klp9 (Figure 6E & F). Moreover, microtubule length, measured just before a growing microtubule underwent catastrophe, increased (Figure 6G) and the catastrophe frequency decreased with increasing Klp9 concentration (Figure 6H). Taken together, the kinesin-6 Klp9 stabilized microtubules and enhanced microtubule growth by increasing the velocity of polymerization and by decreasing the catastrophe frequency. This has similarly been observed for the *S. cerevisiae* plus-end directed kinesin Kip2 (Hibbel et al., 2015) and *X. laevis* kinesin-5 (Chen and Hancock, 2015).

Surprisingly, we found that at a higher tubulin concentration (20 μM), which allowed faster microtubule growth in the control condition (3.21 ± 0.53 μm/min), Klp9 displayed the opposite effect on microtubule growth (Figure 6I & J). With increasing Klp9 concentration the microtubule growth velocity decreased until it reached a plateau, ranging around 2.4 ± 0.5 μm/min in presence of 100 nM Klp9 (Figure 6J). Note that this velocity was similar to the Klp9-mediated microtubule gliding velocity (Figure 6C). Moreover, the microtubule length decreased with increasing Klp9 concentration (Figure 6 – Figure supplement 2) and the catastrophe frequency did not change significantly (Figure 6 – figure supplement 3).

The results suggest that the kinesin-6 does not only promote microtubule polymerization it can also decrease the microtubule growth velocity depending on the tubulin concentration. The kinesin-6 may thus have adopted a mechanism that allows the motor to set a definite microtubule growth velocity, close to its walking speed. To our knowledge, such behavior has not yet been described for kinesins or other microtubule-associated proteins.

## Discussion

This study leads to two main conclusions: (i) The localization of the kinesin-6 Klp9 to spindle microtubules at anaphase onset requires its dephosphorylation at Cdc2-phosphosites, mediated mostly by the XMAP215 family member Dis1 (phosphorylated at Cdc2-phosphosites) and with a smaller proportion by Clp1 (homolog of Cdc14). (ii) Klp9 promotes microtubule polymerization *in vivo* and *in vitro* in a dose-dependent manner. Moreover, *in vitro* at high tubulin concentration, where microtubules grow comparatively fast, an increasing Klp9 concentration caused a decrease of the microtubule growth speed, suggesting that Klp9 acts as a cruise control by setting a distinct microtubule growth velocity.

### Dis1-dependent localization of Klp9 to the anaphase spindle

Members of the XMAP215/Dis1 family have been identified as microtubule-associated proteins that accelerate the rate of microtubule growth from yeast to human (Al-Bassam et al., 2012; Brouhard et al., 2008; Charrasse et al., 1998; Gard and Kirschner, 1987; Kinoshita et al., 2001; Matsuo et al., 2016; Podolski et al., 2014; Tournebize et al., 2000). Moreover, recently XMAP215 family members have been implicated in the regulation of microtubule nucleation (Roostalu et al., 2015; Thawani et al., 2018; Wieczorek et al., 2015). These functions make XMAP215 proteins essential for proper mitotic spindle functioning, with their knockdown leading to small or disorganized spindles (Cassimeris and Morabito, 2004; Cullen et al., 1999; Garcia et al., 2001; Gergely et al., 2003; Goshima et al., 2005; Kronja et al., 2009; Matthews et al., 1998; Ohkura et al., 1988; Reber et al., 2013; Severin et al., 2001; Tournebize et al., 2000). We found that Dis1 is not only important due to its direct effects on spindle microtubules, but also by regulating the recruitment of the mitotic kinesin-6 Klp9. *X. laevis* XMAP215 has been implicated in the recruitment of Cdc2 to spindle microtubules via interaction with Cyclin B (Charrasse et al., 2000), yet a direct effect on mitotic motors has not been reported so far. How Klp9 is dephosphorylated by the phosphorylated form of Dis1 remains elusive. Dis1, phosphorylated at Cdc2-phosphosites, could mediate the activation or release of a phosphatase. It is possible that Dis1, which is still phosphorylated in late metaphase, activates a phosphatase, which subsequently dephosphorylates Klp9 at anaphase onset. Recently, similar to Dis1, the inhibitor of PP2A protein phosphatases, Sds23, has been shown to act upstream of Klp9 and regulate Klp9 recruitment to the spindle (Schutt and Moseley, 2020). Dis1 and Sds23 may act in the same pathway. In general, the results suggest that XMAP215 proteins may also serve as a general regulatory unit of mitotic spindle composition.

Besides, we were wondering about the need for such an unconventional mechanism. Why is Dis1 implicated in the regulation of Klp9 localization to the anaphase spindle? Previously, XMAP215 has been shown to regulate mitotic spindle length (Milunovic-Jevtic et al., 2018; Reber et al., 2013). *In vitro* and *in vivo*, XMAP215 sets spindle length in a dose-dependent manner, and may thus be a crucial factor for the scaling of spindle length to cell size (Krüger and Tran, 2020; Milunovic-Jevtic et al., 2018; Reber et al., 2013). Similarly, Klp9 sets the speed of anaphase B spindle elongation in a dose-dependent manner (Krüger et al., 2019), which allows scaling of anaphase B spindle elongation velocity to spindle length and cell size (Krüger et al., 2019). Thus, the mechanism of Dis1 regulating the recruitment of Klp9 to the anaphase spindle could link the regulation of spindle length to the regulation of spindle elongation velocity: an increased amount of Dis1 molecules results in the formation of longer spindles and the recruitment of more Klp9 molecules, subsequently allowing faster spindle elongation of the longer spindles. To test this hypothesis, we investigated if Dis1 indeed sets spindle length scaling, as it has been suggested by varying the XMAP215 amounts in *X. laevis* cell extracts (Reber et al., 2013) or in *X. laevis* cells of the same size (Milunovic-Jevtic et al., 2018). *Dis1* deletion did not affect spindle length scaling in metaphase (Figure 7 – figure supplement 1), but strongly impacted the scaling relationship in anaphase B (Figure 7A). *Klp9* deletion also diminished the scaling relationship (Figure 7A), either due to its function in microtubule polymerization or sliding, but deletion of *dis1* displayed an even stronger effect, reducing the slope of the linear regression from 0.5 in wild-type cells to 0.21 in *dis1* deleted cells of different sizes (Figure 7A). Thus, Dis1 regulating Klp9 recruitment during anaphase B could indeed link the regulation of spindle length to the regulation of spindle elongation velocity, ensuring that shorter spindles elongate with slower speeds, and longer spindles with higher speeds, as shown previously (Krüger et al., 2019). Eventually, this ensures that longer spindles accomplish chromosome segregation within the same time frame as shorter spindles, thus preventing a prolongation of mitosis, which may be harmful for cell viability (Krüger and Tran, 2020).

**Figure 7:**
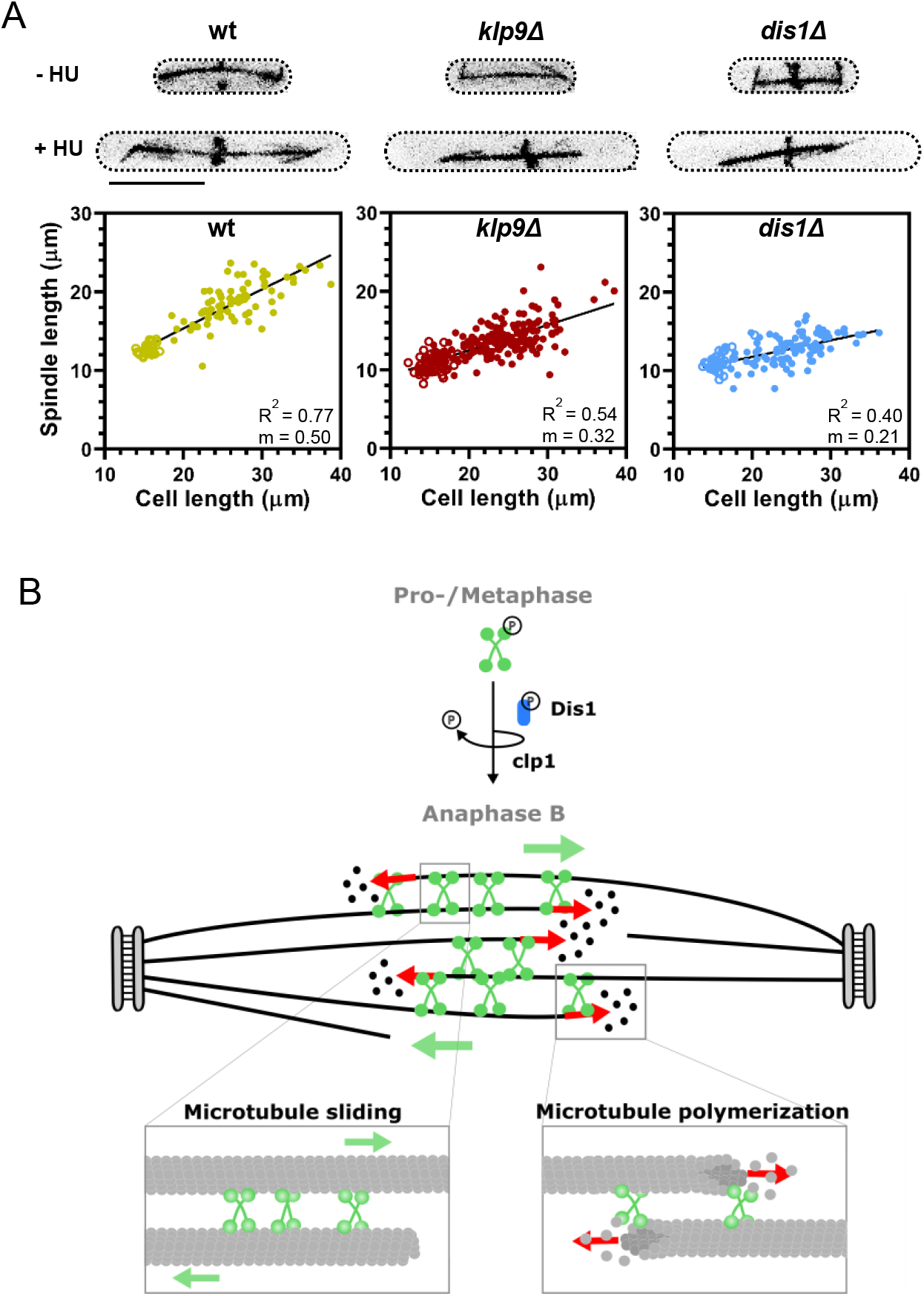
Model of Dis1-dependent Klp9 recruitment and Klp9 function during anaphase B spindle elongation. (A) Upper panel: Wild-type, *klp9Δ* and *dis1Δ* cells at the end of anaphase B expressing mCherry-Atb2 treated or not treated with hydroxyurea (HU). Addition of HU allows to block the cells in S-Phase, resulting in increased cell size. Lower panel: Final anaphase B spindle length plotted against cell length of wild-type, *klp9Δ* and *dis1Δ* cells. Unfilled circles correspond to cells not treated with HU and filled circles to cells treated with HU. Data was fitted by linear regression, showing the regression coefficicent R^2^ and the slope m. (B) Model of Klp9 recruitment at anaphase onset and its function during anaphase B. Klp9 may promote spindle elongation by generating microtubule sliding forces and regulating the microtubule growth velocity.

### Regulation of microtubule dynamics by kinesin-6 Klp9

The *in vivo* and *in vitro* experiments at hand strongly suggest the kinesin-6 Klp9 to be a crucial regulator of microtubule dynamics. In monopolar spindles, in presence of Klp9, microtubule bundles grew with a velocity that matches the microtubule growth velocity expected in bipolar spindles. In absence of Klp9, the growth velocity and length of microtubule bundles was strongly diminished, indicating that the microtubule growth necessary for anaphase B spindle elongation is promoted by the kinesin-6. This is supported by the observation that Klp9 prominently localized to and accumulated at microtubule bundle tips with anaphase B progression. Microtubule plus-end accumulation appears to be an important function for microtubule dynamics regulating kinesins (Chen and Hancock, 2015; Gudimchuk et al., 2013; Hibbel et al., 2015; Varga et al., 2006, 2009). We have not observed such a strong tip-tracking activity for other anaphase B spindle components in monopolar spindles, namely the microtubule bundler Ase1, the microtubule rescue factor CLASP, the XMAP215 protein Dis1, or the other bipolar sliding motor kinesin-5. We thus argue, that during anaphase B, Klp9 is crucial for the regulation of microtubule dynamics.

*In vitro* reconstitution experiments with recombinant full-length Klp9 provide further evidence for this role of the sliding motor. With low microtubule growth rates in the control condition, Klp9 increased the growth velocity in a dose-dependent manner. This has similarly been observed for the *X. laevis* kinesin-5 Eg5 (Chen and Hancock, 2015). The fact that not only a dimeric, but also a monomeric Eg5 construct promoted microtubule polymerization, suggests that the underlying mechanism is based on Eg5 stabilizing a straight tubulin conformation (Chen et al., 2019, 2016; Chen and Hancock, 2015). In solution, tubulin exhibits a curved conformation, which is not potent for microtubule lattice incorporation (Ayaz et al., 2012; Rice et al., 2008; Richard McIntosh et al., 2018). The motor walks toward the microtubule plus-end, accumulates there by staying bound for time periods that greatly exceed the stepping duration (Chen and Hancock, 2015), and straightens newly added tubulin dimers, thus stabilizing lateral tubulin-tubulin interactions and promoting microtubule growth (Chen et al., 2019). This differs from the mechanism of microtubule polymerization proposed for members of the XMAP215 family. In contrast to kinesin-5, XMAP215 proteins show high affinity for free tubulin (Ayaz et al., 2014; Brouhard et al., 2008). XMAP215 TOG domains are proposed to bring tubulin close to the plus ends so that it can be incorporated into the microtubule lattice (Ayaz et al., 2014; Brouhard et al., 2008; Geyer et al., 2018). The mode of action of kinesin-5 and XMAP215 may thus differ temporally: XMAP215 promotes tubulin dimer association with the protofilament and kinesin-5 stabilizes this association.

Interestingly, for kinesin-6 we observed that at higher microtubule growth velocities in the control condition, Klp9 displayed a negative effect on microtubule growth. It decreased the growth velocity in a dose-dependent manner. Due to this observation, we hypothesize that the mechanism of microtubule growth regulation is different from the mechanisms that have been suggested for kinesin-5 (Chen et al., 2019; Chen and Hancock, 2015) or XMAP215 (Ayaz et al., 2014; Brouhard et al., 2008). First, for kinesin-5 and XMAP215 proteins, addition of recombinant protein to higher tubulin concentrations further increases microtubule growth velocities (Al-Bassam et al., 2012; Brouhard et al., 2008; Chen et al., 2019). Second, the fact that microtubule growth velocities converge with addition of Klp9 in presence of low or high tubulin concentrations indicates that Klp9 sets a distinct microtubule growth velocity, and suggests a mechanism beyond a stabilizing effect proposed for kinesin-5 (Chen et al., 2019) or an enhancement of tubulin dimer addition as regulated by XMAP215 (Brouhard et al., 2008). In contrast, to be able to accelerate and decelerate microtubule growth, Klp9 has to promote tubulin dimer addition at microtubule plus ends, as well as to block it. This is reminiscent of the processive elongator of actin filaments formin (Courtemanche, 2018). Formins located at the barbed ends of actin filaments, consisting of a dimer of formin homology 2 domains (FH2), form a ring-like structure around the actin filament, and can accommodate a ‘closed’ conformation blocking actin subunit addition, or an ‘open’ conformation, which allows for subunit addition (Vavylonis et al., 2006). The equilibrium between the ‘closed’ and the ‘open’ conformation of FH2 dimers subsequently determines the rate of actin elongation (Gurel et al., 2015; Kovar et al., 2006; Thompson et al., 2013). Accordingly, one Klp9 molecule located at the end of each protofilament could promote the association of new tubulin dimers with a certain rate, but block the addition beyond that rate. One motorhead could be bound to the very last tubulin dimer, while the other motorhead binds free tubulin and promotes its incorporation into the microtubule lattice. How the inhibition of tubulin subunits occurs appears to be a complicated question. One possibility is that the binding of Klp9 to free curved tubulin induces a conformational change of tubulin dimers that is much more amenable for lattice incorporation. Like kinesin-5, kinesin-6 could promote tubulin dimer straightening, but unlike kinesin-5, not after lattice incorporation, but before. Even if a free tubulin dimer is in close proximity to the last tubulin dimer in the protofilament where motorhead A of Klp9 is bound, the straightened tubulin dimer bound to motorhead B will be more potent for addition to the protofilament than a free curved tubulin dimer. Consequently, the velocity of microtubule growth may be set by the stepping rate of the motor. Accordingly, we found the reduced microtubule growth velocity in presence of Klp9 at high tubulin concentrations to be comparable to the Klp9-mediated microtubule gliding velocity.

Collectively, this work sheds light on the mechanism of Klp9 recruitment and function during anaphase B. Dephosphorylation of Klp9 mediated by Dis1 promotes its localization to the spindle midzone, where it can subsequently promote anaphase B spindle elongation (Figure 7B). This mechanism of Klp9 recruitment may link the regulation of mitotic spindle length and spindle elongation velocity to achieve scaling with cell size. Moreover, we show that anaphase B spindle elongation is achieved not only by Klp9 generating microtubule sliding forces, as shown previously (Fu et al., 2009; Rincon et al., 2017), but also by regulating microtubule polymerization. This makes the kinesin-6 a perfect candidate responsible for the coordination of microtubule sliding and growth (Figure 7B). With Klp9 at the spindle midzone, the microtubule sliding velocity and the microtubule growth velocity of plus-ends located at the edge of the midzone are set with similar speeds. Last, the *in vitro* results suggest an unconventional mechanism of the regulation of microtubule growth. The kinesin-6 appears to not simply enhance tubulin dimer addition at microtubule plus ends, as suggested for other microtubule polymerases (Ayaz et al., 2014; Brouhard et al., 2008; Chen et al., 2019; Chen and Hancock, 2015), but to also block tubulin dimer addition that exceeds a certain rate. Hence, Klp9 may have adopted a mechanism that is comparable to that of formins by being able to promote and block subunit addition to the filament. Eventually, Klp9 can thus set a well-defined microtubule growth velocity of slow and fast growing microtubules within the spindle, and act as a ‘cruise control’ of microtubule polymerization.

## Acknowledgements

We thank Manuel Lera Ramirez and Ana Loncar for discussions and critical reading of the manuscript. We thank Moutse Ranaivoson for help with the Klp9 purification. We thank Vincent Fraisier and Lucy Sengmanivong for the maintenance of microscopes at the PICT-IBiSA Imaging facility (Institut Curie), a member of the France-BioImaging national research infrastructure. We thank the Japan National BioResource Project - Yeast Genetic Resource Center (Osaka City University, Osaka University and Hiroshima University) for providing strains. We thank Chris Norbury (Oxford University) for generously providing the *S. pombe* cDNA library.

## Materials & Methods

### Production of *S. pombe* mutant strains

All used strains are isogenic to wild-type 972 and were obtained from genetic crosses, selected by random spore germination and replica on plates with appropriate drugs or supplements. All strains are listed in the supplementary strain list. Gene deletion and tagging was performed as described previously (Bähler et al., 1998).

### Fission yeast culture

All *S. pombe* strains were maintained at 25°C and cultured in standard media. Strains were either maintained on YE5S plates or EMM plates supplemented with adenine, leucine and uracil in case of the Klp9 Shut-Off strain, at 25°C and refreshed every third day. In general, cells were transferred into liquid YE5S and imaged the following day at exponential growth.

For overexpression of *dis1*, strains were transferred to liquid EMM supplemented with adenine, leucine, uracil and thiamine two days before imaging. The following day cells were centrifuged at 3000 rpm for 5 min, washed three times with H2O and resuspended in EMM supplemented with adenine, leucine and uracil. Following incubation at 25°C for 18-20 hours cells were imaged.

For shut-off of Klp9, cells were cultured in EMM supplemented with adenine, leucine and uracil two days before imaging. The following day, cells were centrifuged at 3000 rpm for 5 min, resuspended in YE5S and incubated at 25°C for approximately 20 hours until imaging.

The generation of long cells was achieved by treatment with Hydroxyurea (Sigma Aldrich). Cells were transferred to liquid YE5S at 25°C and 10 mM 10mM Hydroxyurea was added when cells reached exponential growth. After incubation for 5 hours at 25°C, cells were washed three times with H2O and resuspended in fresh YE5S. Following one hour at 25°C cells were imaged.

### Live microscopy of *S. pombe* cells

For live-cell imaging, fission yeast cells were mounted on YE5S agarose pads, containing 2% agarose (Tran et al., 2004). Temperature-sensitive *cut7-24* and *sad1-1* mutants, were incubated at the microscope, which is equipped with a cage incubator (Life Imaging Services), at 37°C for one hour before imaging was started. All other strains were imaged at 25°C.

Images were acquired on an inverted Eclipse Ti-E microscope (Nikon) with a spinning disk CSU-22 (Yokogawa), equipped with a Plan Apochromat 100x/1.4 NA objective lens (Nikon), a PIFOC (perfect image focus) objective stepper, a Mad City Lab piezo stage and an electron-multiplying charge-couple device camera (EMCCD 512×512 Evolve, Photometrics).

Stacks of 7 planes spaced by 1 μm were acquired for each channel with 100 ms exposure time, binning 1 and an electronic gain of 300 for both wavelengths. For each time-lapse movie an image was taken every minute for 90-120 minutes.

### Construction, expression and purification of recombinant Klp9

For construction of a plasmid containing full-length Klp9, *klp9* cDNA was amplified from a cDNA library (Hoffman et al., 2015) and cloned into pET28a(+) between NcoI and NotI restriction sites, incorporating a C-terminal His6 tag.

For Klp9 purification *E. coli* NiCO21 (DE3) cells were transfected with pET28a-Klp9-His6 and grown in standard 2xYT medium. Following cell lysis and centrifugation, the supernatant was incubated with chitin resin (New England Biolabs) for 45 minutes at 4°C and eluted on a gravity flow column. The protein solution was then loaded on an HiTrap IMAC HP column (GE Healthcare). Further purification of the Klp9-containing fractions was achieved by gel filtration using a Superdex 200 26/60 column equilibrated with a buffer containing 20 mM HEPES pH 8, 150 mM NaCl, 2 mM MgCl2, 0.2 mM ADP, 0.5 mM TCEP, 0.5 mM PMSF and 10% Glycerol.

### Tubulin purification

Tubulin from fresh bovine brain was purified by three cycles of temperature-dependent assembly and disassembly in BRB80 buffer. Labeling of tubulin with ATTO-488, ATTO-647 or biotin was performed as previously described (Hyman et al., 1991).

### *In vitro* microtubule gliding assays

Taxol-stabilized microtubules were prepared by incubating 56 μM unlabeled tubulin and 14 μM ATTO-647-labeled tubulin in BRB80 (80 mM piperazineN,N[prime]-bis(2-ethanesulfonic acid (PIPES) pH 6.8, 1 mM EGTA, 1 mM MgCl2) with 20 mM MgCl2, 10 mM GTP, 25% DMSO in BRB80 at room temperature. The mixture was then diluted 1:111 with 10 μM taxol in BRB80.

The flow chamber was assembled from microscopy slides and glass coverslips cleaned sequentially in acetone in a sonication bath and in ethanol. Then, 0.1 mg/ml Anti-His6 (Sigma Aldrich) was added to the flow chamber, followed by 30 μM recombinant Klp9. Upon subsequent incubation of the flow chamber with taxol-stabilized microtubules, a buffer containing 0.5% Methylcellulose, 125 mM KCl, 12.5 mM MgCl2, 2.5 mM EGTA, 16.6 mM Hepes pH 7.5, 3 mg/ml Glucose, 0.1 mg/ml Glucose Oxydase, 0.03 mg/ml Catalase, 8.5 mM ATP and 0.3% BSA was added and the flow chamber was sealed with vacuum grease. Imaging was performed immediately after at 30°C at the TIRF microscope.

### *In vitro* microtubule dynamics assays

GMPCPP-stabilized microtubule seeds were prepared by incubating a mixture of 7 μM biotinylated tubulin, 3 μM ATTO-647-labeled tubulin and 0.5 mM GMPCPP (Interchim) in 1xBRB80 for one hour at 37°C. Following, 2.5 μM taxol was added and the mix was incubated for 20 minutes at room temperature (RT). After centrifugation at 14000 rpm for 10 minutes the supernatant was removed and the pellet resuspended in a mix containing 2.5 μM taxol and 0.5 μM GMPCPP in 1xBRB80. The seeds were then snap frozen in liquid nitrogen and stored at −80°C.

Flow chambers were assembled from microscopy slides and glass coverslips with double-sticky tape with three independent lanes, allowing the analysis of three different reaction mixtures in the same flow chamber. Slides and coverslips were cleaned sequentially in acetone in a sonication bath, in ethanol, in 2% Hellmanex (Sigma Aldrich) and using a plasma-cleaner. Functionalization was achieved by sequential incubation with 1 mg/ml PEG-Silane (30 K, Creative PEGWorks) for microscopy slides or 0.8 mg/ml PEG-Silane and 0.2 mg/ml Biotin-PEG-Silane (5 K, Creative PEGWorks) for coverslips and 0.05 mg/ml NeutrAvidin (Invitrogen) in G-Buffer (2 mM Tris-Cl pH 8, 0.2 mM ATP, 0.5 mM DTT, 0.1 mM CaCl2, 1 mM NaAzide, 6.7 mM Hepes pH 7.5, 50 mM KCl, 5 mM MgCl2 and 1mM EGTA). Attachment of microtubule seeds to the coverslip was achieved through biotin-neutrAvidin interaction. The reaction mixture with or without Klp9 in G-Buffer contained a tubulin mix (80% unlabeled tubulin, 20% 488-labeled tubulin in BRB80) 0.5% Methylcellulose, 50 mM KCl, 5 mM MgCl2, 1 mM EGTA, 6.6 mM Hepes pH 7.5, 3 mg/ml Glucose, 0.1 mg/ml Glucose Oxydase, 0.03 mg/ml Catalase, 6.6 mM Tris-Cl pH 8, 0.65 mM ATP, 1 mM GTP, 11.5 mM DTT, 0.3 mM CaCl2, 3.3 mM NaAzide., 0.1% BSA. The mixture was added to the flow chamber, which was then sealed with vacuum grease and the dynamic microtubules were imaged immediately at 30°C at the TIRF microscope.

### TIRF microscopy

*In vitro* reconstitution assays were imaged on an inverted Eclipse Ti-E (Nikon) equipped with Perfect Focus System (PFS 2, Nikon), a CFI Plan Apo TIRF 100x/1.49 N.A oil objective (Nikon), an TIRF-E motorized TIRF illuminator and an EMCCD Evolve camera (Photometrics). For excitation 491 nm 42 mW and 642 nm 480 mW (Gataca Systems) were used. The temperature was controlled with a cage incubator (Life Imaging Services).

For microtubule gliding and microtubule dynamics assays, movies of 5 min duration with 1 second interval were acquired.

### Image analysis

Using Metamorph 7.2, maximum projections of each stack were performed for analysis of spindle dynamics and for presentation and sum projections for intensity measurements.

Spindle and bundle dynamics were examined by the length of the mCherry-Atb2 signal over time. Metaphase and anaphase spindle length refer to the final spindle length of each phase.

Intensity measurements were performed by drawing a region around the area of interest, reading out the average intensity per pixel, subtracting the background and multiplying this value with the size of the area. Intensity profiles along microtubule bundles or along the bipolar spindle were obtained by drawing a line along the bundle/spindle and reading out the intensity values for each pixel along the line, from which the background intensity was subtracted.

The microtubule gliding velocity in the *in vitro* assays was revealed by constructing kymographs of the moving microtubules using Metamorph and calculation of the velocity by the microtubule displacement within each 5 min movie.

To analyze microtubule dynamics parameters of individual microtubules growing from GMPCPP-seeds kymographs were cosntructed with Metamorph. For each microtubule a kymograph was constructed and the slope of growth and shrinkage events as well as the microtubule length, corresponding to the length of a microtubule before undergoing catastrophe, was analyzed. The catastrophe frequency was calculated by the number of catastrophe events per microtubule within the 5 minute movie divided by the total time the microtubule spent growing.

### Quantification and statistical analysis

Sample number, replicates and error bars are indicated in the figure legends. All statistical analysis and regression analysis were performed using GraphPad Prism 7. Corresponding details for statistical tests and p values are included in the figures and/or figure legends.

## Figure supplements

**Figure 1 – figure supplement 1:**
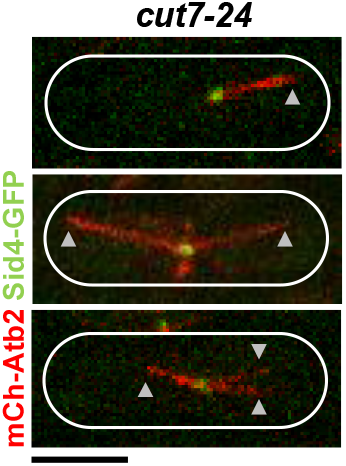
*Cut7-24* cells expressing mCherry-Atb2 (tubulin) and Sid4-GFP (SPBs) at 37°C. Arrowheads depict the tip of microtubule bundles growing from the unseparated spindle poles.

**Figure 3 – figure supplement 1:**
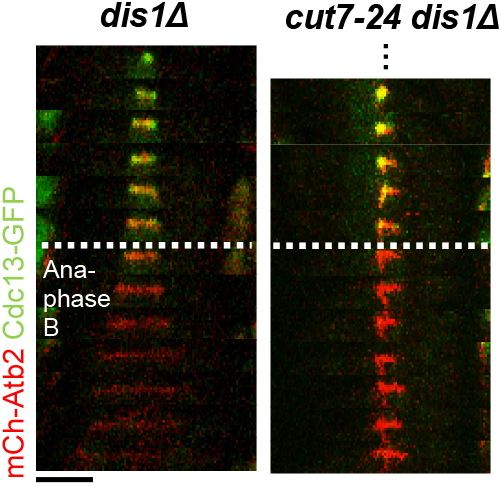
Time-lapse images of *dis1Δ* and *cut7-24* dis1Δ cells expressing mCherry-Atb2 (tubulin) and Cdc13-GFP (Cyclin B) at 37°C. Dotted line denotes the metaphase-to-anaphase B transition. Each frame corresponds to 2 min interval. Scale bar, 5 μm.

**Figure 3 – figure supplement 2:**
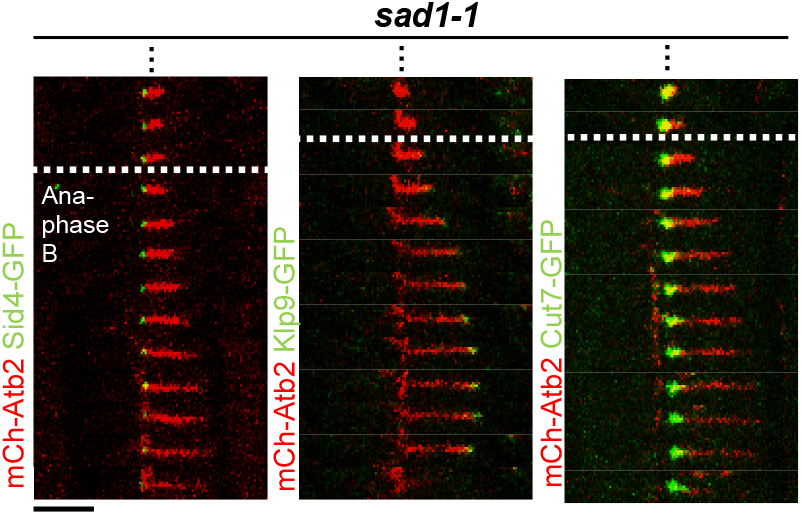
Time-lapse images of *sad1-1* cells expressing mCherry-Atb2 (tubulin) and Sid1-GFP (SPBs) or Klp9-GFP or Cut7-GFP at 37°C. Dotted line denotes the metaphase-to-anaphase B transition. Each frame corresponds to 1 min interval. Scale bar, 5 μm.

**Figure 3 – figure supplement 3:**
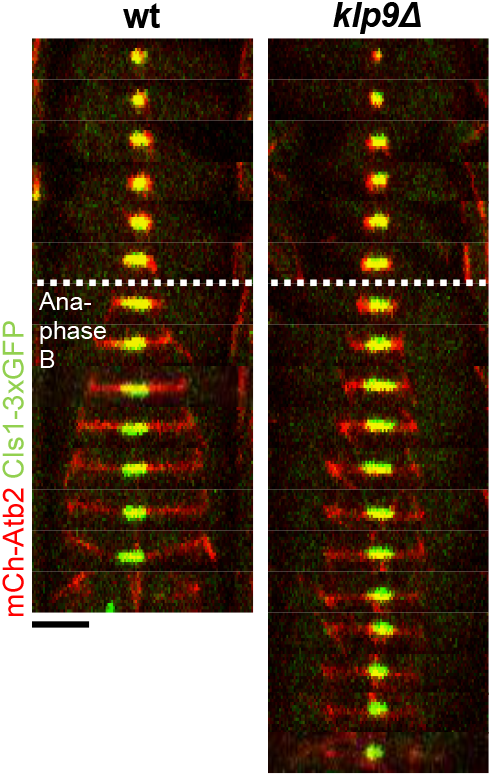
Time-lapse images of wildtype and *klp9Δ* cells expressing mCherry-Atb2 (tubulin) and Cls1-3xGFP (CLASP) at 25°C. Dotted line denotes the metaphase-to-anaphase B transition. Each frame corresponds to 2 min interval. Scale bar, 5 μm.

**Figure 3 – figure supplement 4:**
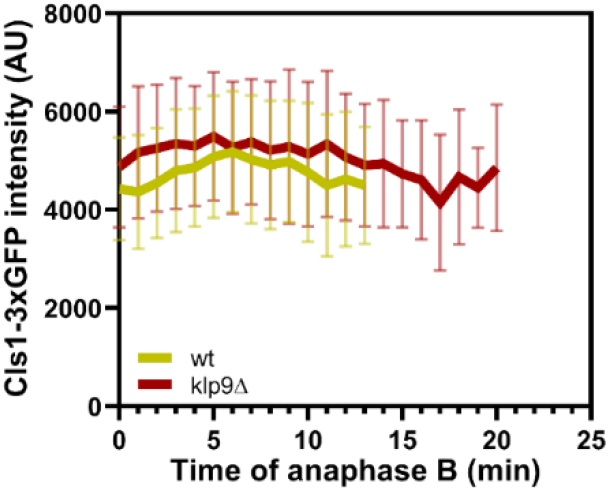
Comparative plot of Cls1-3xGFP intensity throughout anaphase B spindle elongation of wild-type (n=20) and *klp9Δ* (n=20). Bold curves correspond to the mean and error bars to the standard deviation.

**Figure 3 – figure supplement 5:**
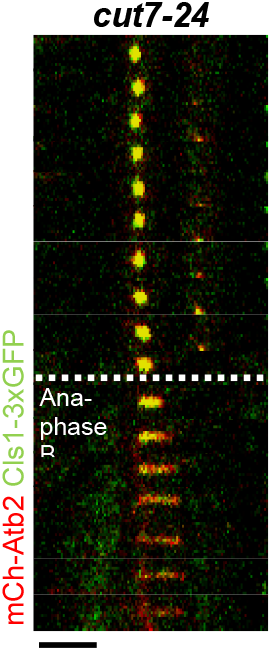
Time-lapse images of *cut7-24* cells expressing mCherry-Atb2 (tubulin) and Cls1-3xGFP (CLASP) at 37°C. Dotted line denotes the metaphase-to-anaphase B transition. Each frame corresponds to 2 min interval. Scale bar, 5 μm.

**Figure 4 – figure supplement 1:**
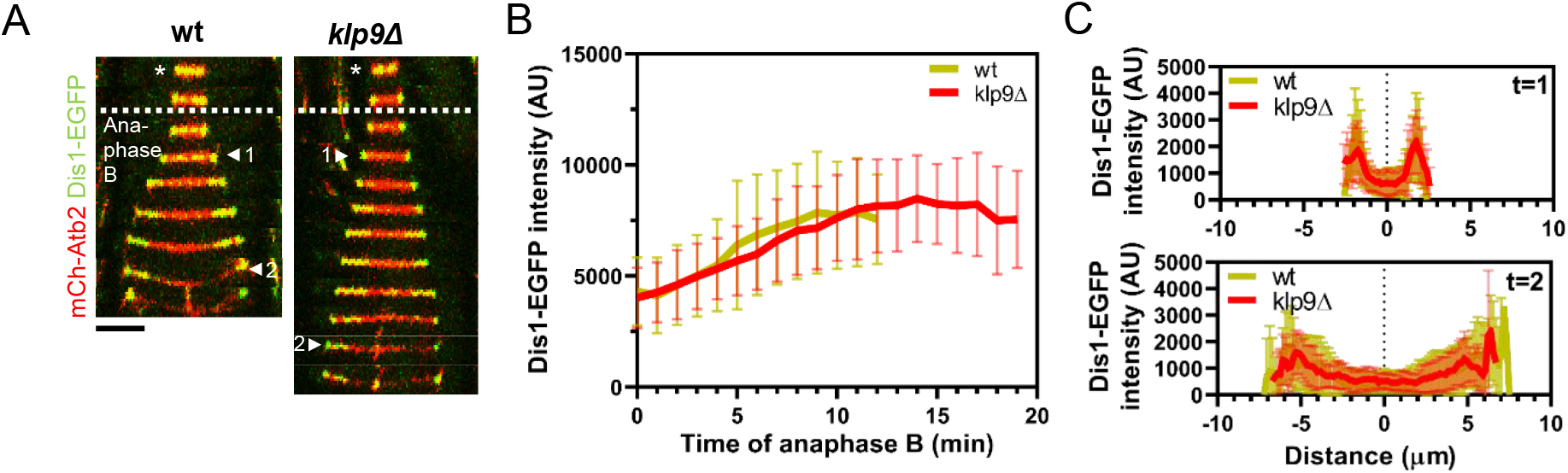
(A) Time-lapse images of wild-type and *klp9Δ* cells expressing mCherry-Atb2 (tubulin) and Dis1-EGFP at 25°C. Each frame corresponds to 2 min interval. Dotted denotes the transition to anaphase B. Arrowheads 1 and 2 correspond to the time points used for linescan analysis at t=1 (2 min after anaphase B onset), t=2 (2 min before spindle disassembly). Scale bar, 5 μm. (B) Comparative plot of Dis1-EGFP intensity throughout anaphase B spindle elongation of wild-type (n=29) and *klp9Δ* (n=29). Bold curves correspond to the mean and error bars to the standard deviation. (C) Intensity spectra obtained by linescan analysis of Dis1-EGFP signals along the anaphase spindle at early (t=1) and late anaphase (t=2). Bold curves correspond to the mean and error bars to the standard deviation.

**Figure 4 – figure supplement 2:**
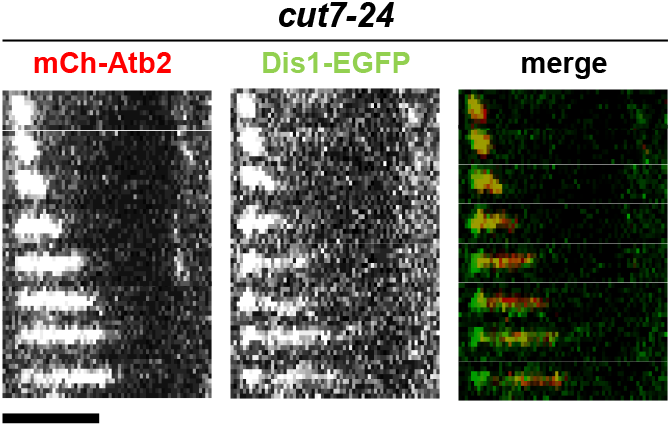
Time-lapse images of *cut7-24* cells expressing mCherry-Atb2 (tubulin) and Dis1-EGFP at 37°C. Each frame corresponds to 2 min interval. Scale bar, 5 μm.

**Figure 4 – figure supplement 3:**
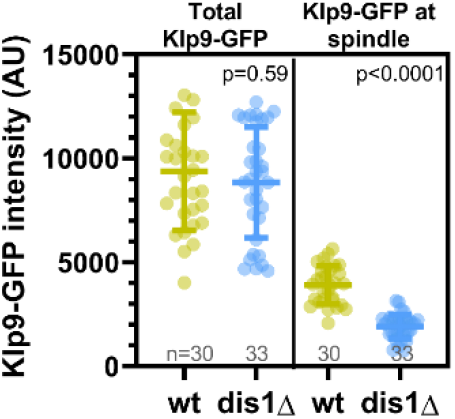
Dot plot comparison of Klp9-GFP intensity in the nucleoplasm at mitosis onset and mean Klp9-GFP intensity at the spindle midzone during anaphase B in wild-type and *dis1Δ* cells. Lines correspond to mean and standard deviation. P values were calculated using Mann-Whitney U test.

**Figure 4 – figure supplement 4:**
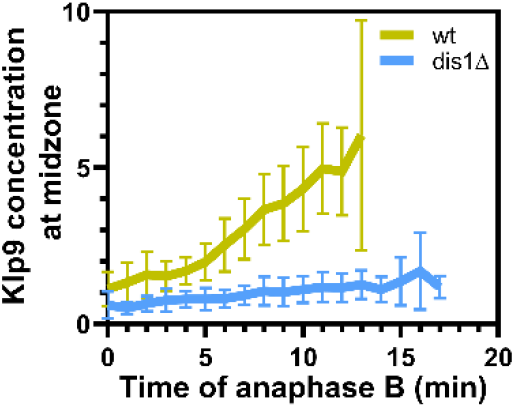
Comparative plot of the relative Klp9 concentration at the spindle midzone (Klp9-GFP devided by mCherry-Atb2 intensity) throughout anaphase B spindle elongation of wildtype (n=29) and *dis1Δ* (n=28). Bold curves correspond to mean spindle dynamics and error bars to the standard deviation.

**Figure 4 – figure supplement 5:**
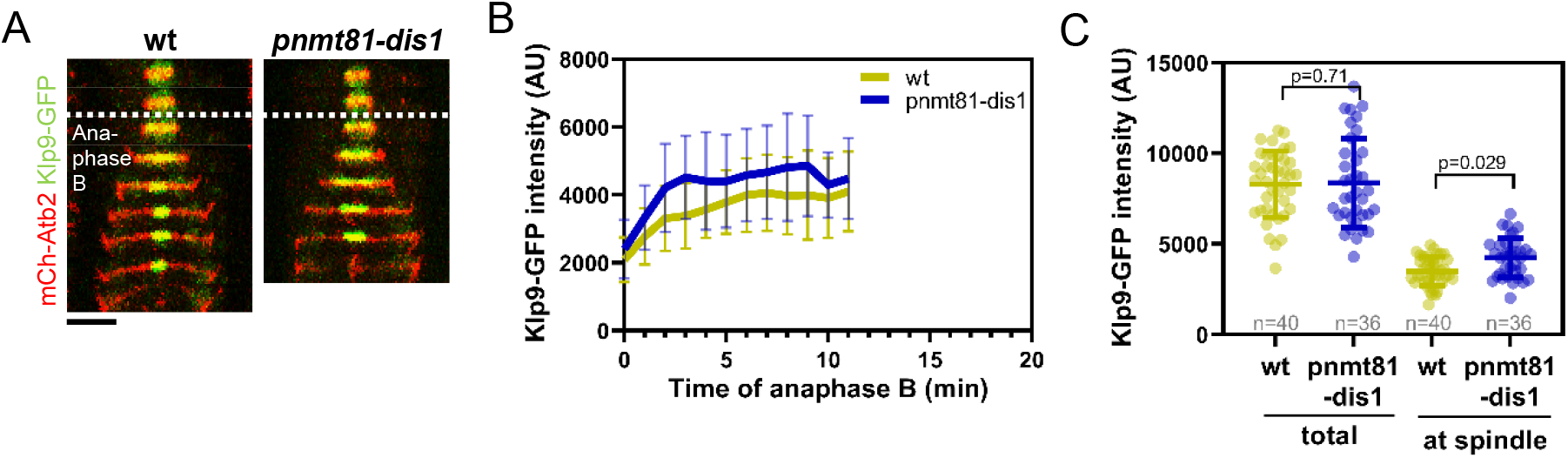
(A) Time-lapse images of wild-type and *pnmt81-dis1* cells expressing mCherry-Atb2 (tubulin) and Klp9-GFP at 25°C. Dotted line corresponds to transition to anaphase B.(B) Comparative plot of Klp9-GFP intensity throughout anaphase B spindle elongation of wild-type (n=40) and *pnmt81-dis1* cells (n=36). Bold curves correspond to the mean and error bars to the standard deviation.(C) Dot plot comparison of Klp9-GFP intensity in wild-type and *pnmt81-dis1* cells in total, measured in the nucleoplasm before anaphase B onset and at the spindle midzone during anaphase B. Lines correspond to mean and standard deviation. P values were calculated using Mann-Whitney U test.

**Figure 5 – figure supplement 1:**
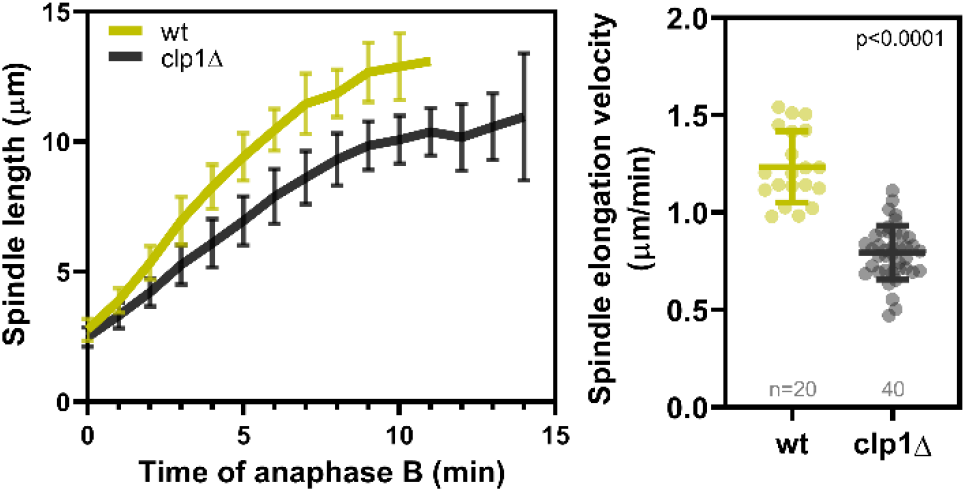
Left panel: Comparative plot of spindle length dynamics during anaphase B of wild-type (n=20) and *clp1Δ* cells (n=40). Bold curves correspond to the mean and error bars to the standard deviation. Right panel: Dot plot comparison of spindle elongation velocity in wild-type and *clp1Δ* cells. Lines correspond to mean and standard deviation. P values were calculated with Mann-Whitney U test.

**Figure 5 – figure supplement 2:**
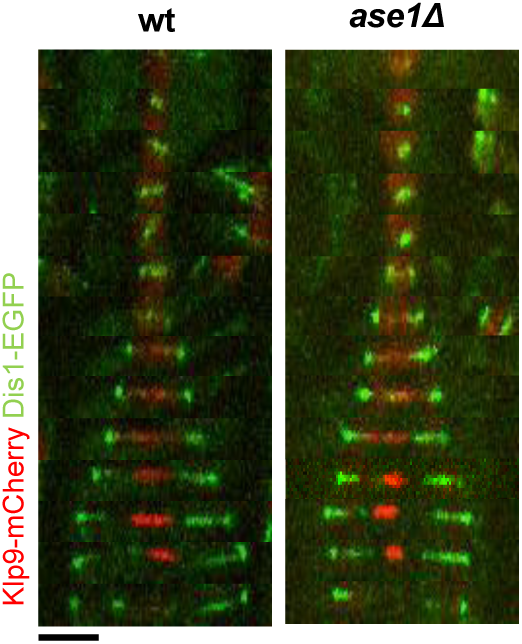
Time-lapse images of wild-type and *ase1Δ* cells expressing Klp9-mCherry and Dis1-EGFP at 25°C. Each time frame corresponds to 2 min interval. Scale bar, 5 μm.

**Figure 5 – figure supplement 3:**
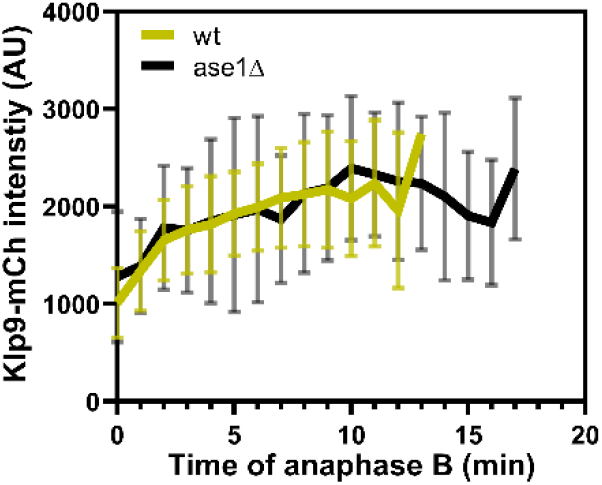
Comparative plot of Klp9-mCherry intensity throughout anaphase B spindle elongation of wild-type (n=30) and *ase1Δ* (n=30). Bold curves correspond to the mean and error bars to the standard deviation.

**Figure 6 – figure supplement 1:**
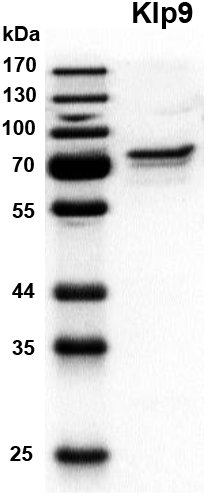
SDS-Gel of purified full-length Klp9 (71 kDa) stained with Instant Blue (Euromedex).

**Figure 6 – figure supplement 2:**
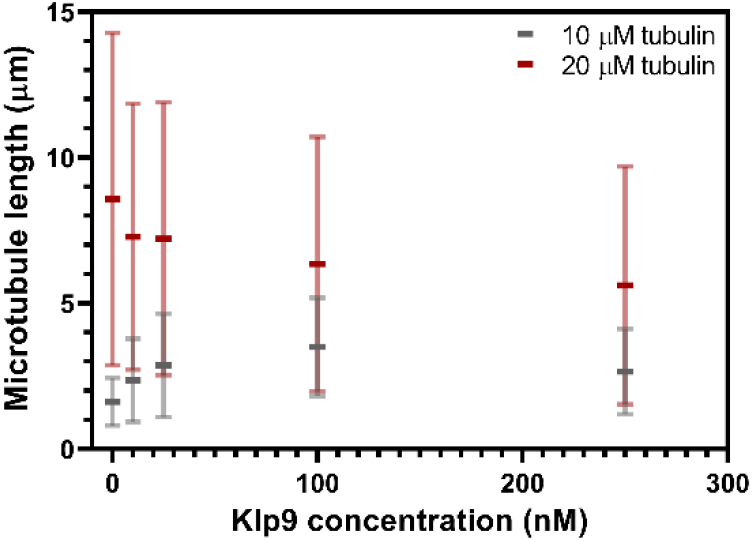
Microtubule length (μm) measured in presence of 10 μM or 20 μM free tubulin shown as a function of the Klp9 concentration (nM); n=190-399 microtubules per condition. Mean values and standard deviations are shown.

**Figure 6 – figure supplement 3:**
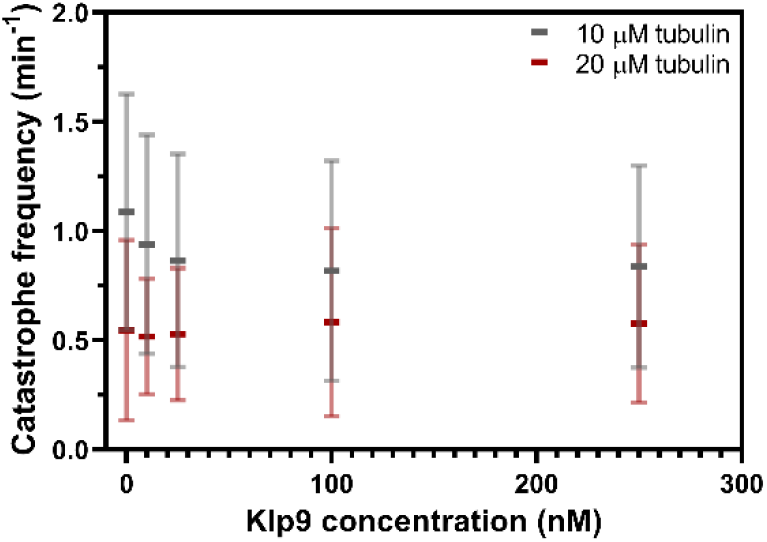
Catastrophe frequency (min^−1^) measured in presence of 10 μM or 20 μM free tubulin shown as a function of the Klp9 concentration (nM); n=190-399 microtubules per condition. Mean values and standard deviations are shown.

**Figure 7 – figure supplement 1:**
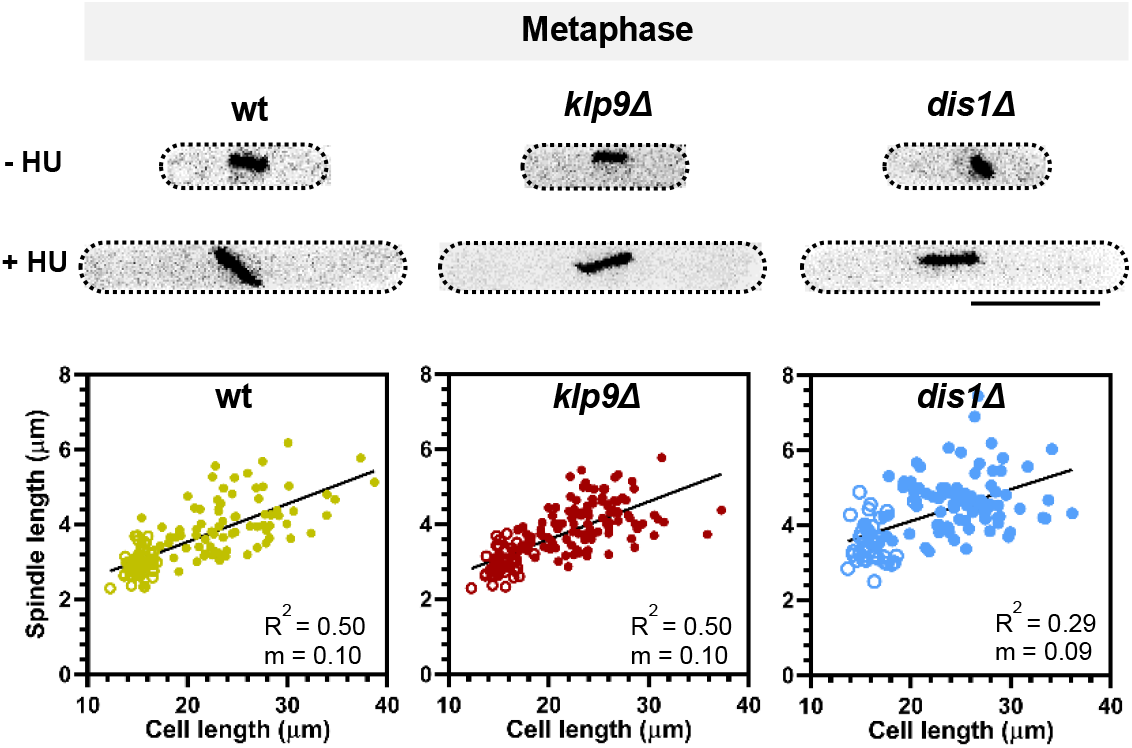
Upper panel: Wild-type, *klp9Δ* and *dis1Δ* cells at the end of metaphase expressing mCherry-Atb2 treared or not treated with hydroxyurea (HU). Addition of HU allows to block the cells in S-Phase, resulting in increased cell size. Lower panel: Final metaphase spindle length plotted against cell length of wild-type, *klp9Δ* and *dis1Δ* cells. Unfilled circles correspond to cells not treated with HU, filled circles to cells treated with HU. Data was fitted by linear regression, showing the regression coefficicent R^2^ and the slope m.

## References

Aist JR, Liang H, Berns MW. 1993. Astral and spindle forces in PtK 2 cells during anaphase B: a laser microbeam study. J Cell Sci 104:1207–1216.

Al-Bassam J, Kim H, Flor-Parra I, Lal N, Velji H, Chang F. 2012. Fission yeast Alp14 is a dose-dependent plus end-tracking microtubule polymerase. Mol Biol Cell 23:2878–2890. doi:10.1091/mbc.E12-03-0205

Aoki K, Nakaseko Y, Kinoshita K, Goshima G, Yanagida M. 2006. Cdc2 Phosphorylation of the Fission Yeast Dis1 Ensures Accurate Chromosome Segregation. Curr Biol 16:1627–1635. doi:10.1016/j.cub.2006.06.065

Avunie-Masala R, Movshovich N, Nissenkorn Y, Gerson-Gurwitz A, Fridman V, Koivomagi M, Loog M, Hoyt MA, Zaritsky A, Gheber L. 2011. Phospho-regulation of kinesin-5 during anaphase spindle elongation. J Cell Sci 124:873–878. doi:10.1242/jcs.077396

Ayaz P, Munyoki S, Geyer EA, Piedra FA, Vu ES, Bromberg R, Otwinowski Z, Grishin N V., Brautigam CA, Rice LM. 2014. A tethered delivery mechanism explains the catalytic action of a microtubule polymerase. Elife 3:1–19. doi:10.7554/eLife.03069

Ayaz P, Ye X, Huddelston P, Brautigam CA, Rice LM. 2012. A TOG:ab-tubulin Complex Structure Reveals Conformation-Based Mechanisms for a Microtubule Polymerase. Science 337:857–860. doi:10.1126/science.1221698

Bähler J, Wu JQ, Longtine MS, Shah NG, McKenzie A, Steever AB, Wach A, Philippsen P, Pringle JR. 1998. Heterologous modules for efficient and versatile PCR-based gene targeting in Schizosaccharomyces pombe. Yeast 14:943–951. doi:10.1002/(SICI)1097-0061(199807)14:10<943::AID-YEA292>3.0.CO;2-Y

Bieling P, Laan L, Schek H, Munteanu EL, Sandblad L, Dogterom M, Brunner D, Surrey T. 2007. Reconstitution of a microtubule plus-end tracking system in vitro. Nature 450:1100–1105. doi:10.1038/nature06386

Bratman S V., Chang F. 2007. Stabilization of Overlapping Microtubules by Fission Yeast CLASP. Dev Cell 13:812–827. doi:10.1016/j.devcel.2007.10.015

Brouhard GJ, Stear JH, Noetzel TL, Al-Bassam J, Kinoshita K, Harrison SC, Howard J, Hyman AA. 2008. XMAP215 Is a Processive Microtubule Polymerase. Cell 132:79–88. doi:10.1016/j.cell.2007.11.043

Brust-Mascher I, Sommi P, Scholey DKCJM. 2009. Kinesin-5–dependent Poleward Flux and Spindle Length Control in Drosophila Embryo Mitosis. Mol Biol Cell 20:1749–1762. doi:https://doi.org/10.1091/mbc.e08-10-1033

Cande WZ, Mcdonald K. 1986. Physiological and Ultrastructural Analysis of Elongating Mitotic Spindles Reactivated In Vitro. J Cell Biol 103:593–604. doi:10.1083/jcb.103.2.593

Cande WZ, McDonald KL. 1985. In vitro reactivation of anaphase spindle elongation using isolated diatom spindles. Lett to Nat 316:168–170.

Cassimeris L, Morabito J. 2004. TOGp, the Human Homolog of XMAP215/Dis1, Is Required for Centrosome Integrity, Spindle Pole Organization, and Bipolar Spindle Assembly. Mol Biol Cell 15:1580–1590. doi:10.1091/mbc.E03

Charrasse S, Lorca T, Dorée M, Larroque C. 2000. The Xenopus XMAP215 and its human homologue TOG proteins interact with cyclin B1 to target p34cdc2 to microtubules during mitosis. Exp Cell Res 254:249–256. doi:10.1006/excr.1999.4740

Charrasse S, Schroeder M, Gauthier-Rouviere C, Ango F, Cassimeris L, Gard DL, Larroque C. 1998. The TOGp protein is a new human microtubule-associated protein homologous to the Xenopus XMAP215. J Cell Sci 111:1371–1383.

Cheerambathur DK, Civelekoglu-Scholey G, Brust-Mascher I, Sommi P, Mogilner A, Scholey JM. 2007. Quantitative analysis of an anaphase B switch: Predicted role for a microtubule catastrophe gradient. J Cell Biol 177:995–1004. doi:10.1083/jcb.200611113

Chen GY, Cleary JM, Asenjo AB, Chen Y, Mascaro JA, Arginteanu DFJ, Sosa H, Hancock WO. 2019. Kinesin-5 Promotes Microtubule Nucleation and Assembly by Stabilizing a Lattice-Competent Conformation of Tubulin. Curr Biol 29:2259–2269. doi:10.1016/j.cub.2019.05.075

Chen GY, Mickolajczyk KJ, Hancock WO. 2016. The kinesin-5 chemomechanical cycle is dominated by a two-heads-bound state. J Biol Chem 291:20283–20294. doi:10.1074/jbc.M116.730697

Chen Y, Hancock WO. 2015. Kinesin-5 is a microtubule polymerase. Nat Commun 6:1–10. doi:10.1038/ncomms9160

Costa J, Fu C, Syrovatkina V, Tran PT. 2013. Imaging individual spindle microtubule dynamics in fission yeast. Methods Cell Biol 115:385–394. doi:10.1016/B978-0-12-407757-7.00024-4

Courtemanche N. 2018. Mechanisms of formin-mediated actin assembly and dynamics. Biophys Rev 10:1553–1569. doi:10.1007/s12551-018-0468-6

Cullen CF, Deák P, Glover DM, Ohkura H. 1999. mini spindles: A gene encoding a conserved microtubule-associated protein required for the integrity of the mitotic spindle in Drosophila. J Cell Biol 146:1005–1018. doi:10.1083/jcb.146.5.1005

Decottignies A, Zarzov P, Nurse P. 2001. In vivo localisation of fission yeast cyclin-dependent kinase cdc2p and cyclin B cdc13p during mitosis and meiosis. J Cell Sci 114:2627–2640.

Des Georges A, Katsuki M, Drummond DR, Osei M, Cross RA, Amos LA. 2008. Mal3, the Schizosaccharomyces pombe homolog of EB1, changes the microtubule lattice. Nat Struct Mol Biol 15:1102–1108. doi:10.1038/nsmb.1482

Ding R, McDonald KL, McIntosh JR. 1993. Three-Dimensional Reconstruction and Analysis of Mitotic Spindles from the Yeast, Schizosaccharomyces pombe. J Cell Biol 120:141–151. doi:10.1083/jcb.120.1.141

Euteneuer U, Jackson WT, Mclntosh JR. 1982. Polarity of Spindle Microtubules in Haemanthus Endosperm. J Cell Biol 94:644–653.

Fink G, Schuchardt I, Colombelli J, Stelzer E, Steinberg G. 2006. Dynein-mediated pulling forces drive rapid mitotic spindle elongation in Ustilago maydis. EMBO J 25:4897–4908. doi:10.1038/sj.emboj.7601354

Fu C, Ward JJ, Loiodice I, Velve-Casquillas G, Nedelec FJ, Tran PT. 2009. Phospho-Regulated Interaction between Kinesin-6 Klp9p and Microtubule Bundler Ase1p Promotes Spindle Elongation. Dev Cell 17:257–267. doi:10.1016/j.devcel.2009.06.012

Garcia MA, Vardy L, Koonrugsa N, Toda T. 2001. Fission yeast ch-TOG/XMAP215 homologue Alp14 connects mitotic spindles with the kinetochore and is a component of the Mad2-dependent spindle checkpoint. EMBO J 20:3389–3401. doi:10.1093/emboj/20.13.3389

Gard DL, Kirschner MW. 1987. A microtubule-associated protein from Xenopus eggs that specifically promotes assembly at the plus-end. J Cell Biol 105:2203–2215. doi:10.1083/jcb.105.5.2203

Gergely F, Draviam VM, Raff JW. 2003. The ch-TOG/XMAP215 protein is essential for spindle pole organization in human somatic cells. Genes Dev 17:336–341. doi:10.1101/gad.245603

Geyer EA, Miller MP, Brautigam CA, Biggins S, Rice LM. 2018. Design principles of a microtubule polymerase. Elife 7:1–23. doi:10.7554/eLife.34574

Goshima G, Wollman R, Stuurman N, Scholey JM, Vale RD. 2005. Length control of the metaphase spindle. Curr Biol 15:1979–1988. doi:10.1016/j.cub.2005.09.054

Grill SW, Go P, Hyman AA. 2001. Polarity controls forces governing asymmetric spindle positioning in the Caenorhabditis elegans embryo. Lett to Nat 409:630–633.

Gudimchuk N, Vitre B, Kim Y, Kiyatkin A, Cleveland DW, Ataullakhanov FI, Grishchuk EL. 2013. Kinetochore kinesin CENP-E is a processive bi-directional tracker of dynamic microtubule tips. Nat Cell Biol 15:1079–1088. doi:10.1038/ncb2831

Gurel PS, Mu A, Guo B, Shu R, Mierke DF, Higgs HN. 2015. Assembly and turnover of short actin filaments by the formin INF2 and profilin. J Biol Chem 290:22494–22506. doi:10.1074/jbc.M115.670166

Hagan I, Yanagida M. 1995. The product of the spindle formation gene sad1+ associates with the fission yeast spindle pole body and is essential for viability. J Cell Biol 129:1033–1047. doi:10.1083/jcb.129.4.1033

Hagan IM, Yanagida M. 1992. Kinesin-related cut7 protein associates with mitotic and meiotic spindles in fission yeast. Nature 356:74–76.

Hagan IM, Yanagida M. 1990. Novel potential mitotic motor protein encoded by the fission yeast cut7+ gene. Nature 347:563–566.

Hibbel A, Bogdanova A, Mahamdeh M, Jannasch A, Storch M, Schäffer E, Liakopoulos D, Howard J. 2015. Kinesin Kip2 enhances microtubule growth in vitro through length-dependent feedback on polymerization and catastrophe. Elife 4:1–11. doi:10.7554/eLife.10542

Hoffman CS, Wood V, Fantes PA. 2015. An ancient yeast for young geneticists: A primer on the Schizosaccharomyces pombe model system. Genetics 201:403–423. doi:10.1534/genetics.115.181503

Hu CK, Coughlin M, Field CM, Mitchison TJ. 2011. KIF4 regulates midzone length during cytokinesis. Curr Biol 21:815–824. doi:10.1016/j.cub.2011.04.019

Hussmann F, Drummond DR, Peet DR, Martin DS, Cross RA. 2016. Alp7 / TACC-Alp14 / TOG generates long-lived, fast-growing MTs by an unconventional mechanism. Sci Rep 6:1–15. doi:10.1038/srep20653

Hyman A, Drechsel D, Kellogg D, Salser S, Sawin K, Steffen P, Wordeman L, Mitchison T. 1991. Preparation of modified tubulins. Methods Enzymol 196:478–485. doi:10.1016/0076-6879(91)96041-O

Janson ME, Loughlin R, Loïodice I, Fu C, Brunner D, Nédélec FJ, Tran PT. 2007. Crosslinkers and Motors Organize Dynamic Microtubules to Form Stable Bipolar Arrays in Fission Yeast. Cell 128:357–368. doi:10.1016/j.cell.2006.12.030

Kapitein LC, Peterman EJG, Kwok BH, Kim JH, Kapoor TM, Schmidt CF. 2005. The bipolar mitotic kinesin Eg5 moves on both microtubules that it crosslinks. Lett to Nat 435:114–118. doi:10.1038/nature03493.Published

Katsuki M, Drummond DR, Osei M, Cross RA. 2009. Mal3 masks catastrophe events in Schizosaccharomyces pombe microtubules by inhibiting shrinkage and promoting rescue. J Biol Chem 284:29246–29250. doi:10.1074/jbc.C109.052159

Khodjakov A, La Terra S, Chang F. 2004. Laser Microsurgery in Fission Yeast: Role of the Mitotic Spindle Midzone in Anaphase B. Curr Biol 14:1330–1340. doi:10.1016/j.cub.2004.07.028

Kinoshita K, Arnal I, Desai A, Drechsel DN, Hyman AA. 2001. Reconstitution of physiological microtubule dynamics using purified components. Science (80-) 294:1340–1343. doi:10.1126/science.1064629

Kinoshita K, Noetzel TL, Pelletier L, Mechtler K, Drechsel DN, Schwager A, Lee M, Raff JW, Hyman AA. 2005. Aurora A phosphorylation of TACC3/maskin is required for centrosome-dependent microtubule assembly in mitosis. J Cell Biol 170:1047–1055. doi:10.1083/jcb.200503023

Kiyomitsu T, Cheeseman IM. 2013. Cortical Dynein and Asymmetric Membrane Elongation Coordinately Position the Spindle in Anaphase. Cell 154:391–402. doi:10.1016/j.cell.2013.06.010

Kovar DR, Harris ES, Mahaffy R, Higgs HN, Pollard TD. 2006. Control of the assembly of ATP- and ADP-actin by formins and profilin. Cell 124:423–435. doi:10.1016/j.cell.2005.11.038

Kronja I, Kruljac-Letunic A, Caudron-Herger M, Bieling P, Karsenti E. 2009. XMAP215-EB1 Interaction Is Required for Proper Spindle Assembly and Chromosome Segregation in Xenopus Egg Extract. Mol Biol Cell 20:2673–2683. doi:10.1091/mbc.E08

Krüger LK, Sanchez JL, Paoletti A, Tran PT. 2019. Kinesin-6 regulates cell-size-dependent spindle elongation velocity to keep mitosis duration constant in fission yeast. Elife 8:e42182:1–22. doi:10.7554/eLife.42182

Krüger LK, Tran PT. 2020. Spindle scaling mechanisms. Essays Biochem 64:383–396. doi:10.1042/EBC20190064

Lioutas A, Vernos I. 2013. Aurora A kinase and its substrate TACC3 are required for central spindle assembly. EMBO Rep 14:829–836. doi:10.1038/embor.2013.109

Loiodice I, Staub J, Setty TG, Nguyen N-PT, Paoletti A, Tran PT. 2005. Ase1p Organizes Antiparallel Microtubule Arrays during Interphase and Mitosis in Fission Yeast. Mol Biol Cell 16:1756–1768. doi:10.1091/mbc.E04

Mallavarapu A, Sawin K, Mitchison T. 1999. A switch in microtubule dynamics at the onset of anaphase B in the mitotic spindle of Schizosaccharomyces pombe. Curr Biol 9:1423–1428. doi:10.1016/S0960-9822(00)80090-1

Mastronarde DN, McDonald KL, Ding R, McIntosh JR. 1993. Interpolar spindle microtubules in PTK cells. J Cell Biol 123:1475–1489. doi:10.1083/jcb.123.6.1475

Masuda H. 1995. The formation and functioning of yeast mitotic spindles. BioEssays 17:45–51. doi:10.1002/bies.950170110

Masuda H, Zacheus W. 1987. The Role of Tubulin Polymerization during Spindle Elongation In Vitro. Cell 49:193–202. doi:https://doi.org/10.1016/0092-8674(87)90560-5

Maton G, Edwards F, Lacroix B, Stefanutti M, Laband K, Lieury T, Kim T, Espeut J, Canman JC, Dumont J. 2015. Kinetochore components are required for central spindle assembly. Nat Cell Biol 17:697–705. doi:10.1038/ncb3150

Matsuo Y, Maurer SP, Yukawa M, Zakian S, Singleton MR, Surrey T, Toda T. 2016. An unconventional interaction between Dis1/TOG and Mal3/EB1 promotes the fidelity of chromosome segregation. J Cell Sci 129:4592–4606. doi:10.1242/jcs.197533

Matthews LR, Carter P, Thierry-mieg D, Kemphues K. 1998. ZYG-9, a Caenorhabditis elegans protein required for microtubule organization and function, is a component of meiotic and mitotic spindle poles. J Biol Chem 141:1159–1168.

Mcdonald K, Pickett-heaps JD, Mcintosh JR, Tippit DH. 1977. On the mechanism of anaphase spindle elongation in diatom vulgare. J Cell Biol 74:377–388.

Mcintosh JR, Landis SC. 1971. The distribution of spindle microtubules during mitosis in cultured humand cells. J Cell Biol 49:468–497.

Meadows JC, Lancaster TC, Buttrick GJ, Sochaj AM, Messin LJ, del Mar Mora-Santos M, Hardwick KG, Millar JBA. 2017. Identification of a Sgo2-Dependent but Mad2-Independent Pathway Controlling Anaphase Onset in Fission Yeast. Cell Rep 18:1422–1433. doi:10.1016/j.celrep.2017.01.032

Milunovic-Jevtic A, Jevtic P, Levy DL, Gatlin JC. 2018. In vivo mitotic spindle scaling can be modulated by changing the levels of a single protein: the microtubule polymerase XMAP215. Mol Biol Cell 29:1311–1317. doi:10.1091/mbc.E18-01-0011

Nabeshima K, Kurooka H, Takeuchi M, Kinoshita K, Nakaseko Y, Yanagida M. 1995. p93dis1, which is required for sister chromatid separation, is a novel microtubule and spindle pole body-associating protein phosphorylated at the Cdc2 target sites. Genes Dev 9:1572–1585. doi:10.1101/gad.9.13.1572

Nabeshima K, Nakagawa T, Straight AF, Murray A, Chikashige Y, Yamashita YM, Hiraoka Y, Yanagida M. 1998. Dynamics of Centromeres during Metaphase-Anaphase Transition in Fission Yeast: Dis1 Is Implicated in Force Balance in Metaphase Bipolar Spindle. Mol Biol Cell 9:3211–3225. doi: 10.1091/mbc.9.11.3211

Oegema K, Desai A, Rybina S, Kirkham M, Hyman AA. 2001. Functional analysis of kinetochore assembly in Caenorhabditis elegans. J Cell Biol 153:1209–1225. doi:10.1083/jcb.153.6.1209

Ohkura H, Adachi Y, Kinoshita N, Niwa O, Toda T, Yanagida M. 1988. Cold-sensitive and caffeine-supersensitive mutants of the Schizosaccharomyces pombe dis genes implicated in sister chromatid separation during mitosis. EMBO J 7:1465–1473. doi:10.1002/j.1460-2075.1988.tb02964.x

Olmsted ZT, Colliver AG, Riehlman TD, Paluh JL. 2014. Kinesin-14 and kinesin-5 antagonistically regulate microtubule nucleation by γ-TuRC in yeast and human cells. Nat Commun 5:1–15. doi:10.1038/ncomms6339

Pereira AL, Pereira AJ, Maia ARR, Drabek K, Sayas CL, Hergert PJ, Lince-Faria M, Matos I, Duque C, Stepanova T, Rieder CL, Earnshaw WC, Galjart N, Maiato H. 2006. Mammalian CLASP1 and CLASP2 Cooperate to Ensure Mitotic Fidelity by Regulating Spindle and Kinetochore Function. Mol Biol Cell 17:4526–4542. doi:10.1091/mbc.E06

Peset I, Seiler J, Sardon T, Bejarano LA, Rybina S, Vernos I. 2005. Function and regulation of Maskin, a TACC family protein, in microtubule growth during mitosis. J Cell Biol 170:1057–1066. doi:10.1083/jcb.200504037

Podolski M, Mahamdeh M, Howard J. 2014. Stu2, the Budding Yeast XMAP215 / Dis1 Homolog, Promotes Assembly of Yeast Microtubules by Increasing Growth Rate and Decreasing Catastrophe Frequency. J Biol Chem 289:28087–28093. doi:10.1074/jbc.M114.584300

Reber SB, Baumgart J, Widlund PO, Pozniakovsky A, Howard J, Hyman AA, Jülicher F. 2013. XMAP215 activity sets spindle length by controlling the total mass of spindle microtubules. Nat Cell Biol 15:1116–1122. doi:10.1038/ncb2834

Redemann S, Baumgart J, Lindow N, Shelley M, Nazockdast E, Kratz A, Prohaska S, Brugués J, Fürthauer S, Müller-Reichert T. 2017. C. elegans chromosomes connect to centrosomes by anchoring into the spindle network. Nat Commun 8:1–13. doi:10.1038/ncomms15288

Rice LM, Montabana EA, Agard DA. 2008. The lattice as allosteric effector: Structural studies of αβ-and γ-tubulin clarify the role of GTP in microtubule assembly. PNAS 105:5378–5383. doi:10.1073/pnas.0801155105

Richard McIntosh J, O’Toole E, Morgan G, Austin J, Ulyanov E, Ataullakhanov F, Gudimchuk N. 2018. Microtubules grow by the addition of bent guanosine triphosphate tubulin to the tips of curved protofilaments. J Cell Biol 217:2691–2708. doi:10.1083/jcb.201802138

Rincon SA, Lamson A, Blackwell R, Syrovatkina V, Fraisier V, Paoletti A, Betterton MD, Tran PT. 2017. Kinesin-5-independent mitotic spindle assembly requires the antiparallel microtubule crosslinker Ase1 in fission yeast. Nat Commun 8:1–12. doi:10.1038/ncomms15286

Roostalu J, Cade NI, Surrey T. 2015. Complementary activities of TPX2 and chTOG constitute an efficient importin-regulated microtubule nucleation module. Nat Cell Biol 17:1422–1434. doi:10.1038/ncb3241

Roostalu J, Schiebel E, Khmelinskii A. 2010. Cell cycle control of spindle elongation. Cell Cycle 9:1084–1090. doi:10.4161/cc.9.6.11017

Roque H, Ward JJ, Murrells L, Brunner D, Antony C. 2010. The fission yeast XMAP215 homolog dis1p is involved in microtubule bundle organization. PLoS One 5:1–12. doi:10.1371/journal.pone.0014201

Sato M, Vardy L, Garcia MA, Koonrugsa N, Toda T. 2004. Interdependency of Fission Yeast Alp14/TOG and Coiled Coil Protein Alp7 in Microtubule Localization and Bipolar Spindle Formation. Mol Biol Cell 15:1609–1622. doi:10.1091/mbc.E03

Saunders W, Koshland D, Eshel D, Gibbons I, Hoyt A. 1995. Saccharomycescerevisiae kinesin- and dynein-related protein required for anaphase chromosome segregation. J Cell Biol 128:617–624. doi:10.1083/jcb.128.4.617

Saxton WM, Mclntosh JR. 1987. Interzone Microtubule Behavior in Late Anaphase and Telophase Spindles. J Cell Biol 105:875–886. doi:10.1083/jcb.105.2.875

Scholey J, Civelekoglu-Scholey G, Brust-Mascher I. 2016. Anaphase B. Biology (Basel) 5:1–30. doi:10.3390/biology5040051

Schutt KL, Moseley JB. 2020. The phosphatase inhibitor Sds23 promotes symmetric spindle positioning in fission yeast. Cytoskeleton 1–14. doi:10.1002/cm.21648

Severin F, Habermann B, Huffaker T, Hyman T. 2001. Stu2 Promotes Mitotic Spindle Elongation in Anaphase. J Cell Biol 153:435–442.

Sharp DJ, Brown HM, Kwon M, Rogers GC, Holland G, Scholey JM. 2000. Functional coordination of three mitotic motors in Drosophila embryos. Mol Biol Cell 11:241–253. doi:10.1091/mbc.11.1.241

Sharp DJ, Yu KR, Sisson JC, Sullivan W, Scholey JM. 1999. Antagonistic microtubule-sliding motors position mitotic centrosomes in Drosopnifa early embryos. Nat Cell Biol 1:51–54. doi:10.1038/9025

Shimamoto Y, Forth S, Kapoor TM. 2015. Measuring Pushing and Braking Forces Generated by Ensembles of Kinesin-5 Crosslinking Two Microtubules. Dev Cell 34:669–681. doi:10.1016/j.devcel.2015.08.017

Straight AF, Sedat JW, Murray AW. 1998. Time-lapse microscopy reveals unique roles for kinesins during anaphase in budding yeast. J Cell Biol 143:687–694. doi:10.1083/jcb.143.3.687

Syrovatkina V, Tran PT. 2015. Loss of kinesin-14 results in aneuploidy via kinesin-5-dependent microtubule protrusions leading to chromosome cut. Nat Commun 6:1–8. doi:10.1038/ncomms8322

Thawani A, Kadzik RS, Petry S. 2018. XMAP215 is a microtubule nucleation factor that functions synergistically with the γ-tubulin ring complex. Nat Cell Biol 20:575–585. doi:10.1038/s41556-018-0091-6

Thompson ME, Heimsath EG, Gauvin TJ, Higgs HN, Jon Kull F. 2013. FMNL3 FH2-actin structure gives insight into formin-mediated actin nucleation and elongation. Nat Struct Mol Biol 20:111–118. doi:10.1038/nsmb.2462

Tolic-Norrelykke I, Sacconi L, Thon G, Pavone FS. 2004. Positioning and Elongation of the Fission Yeast Spindle by Microtubule-Based Pushing. Curr Biol 14:1181–1186. doi:10.1016/j.cub.2004.06.029

Tournebize R, Popov A, Kinoshita K, Ashford AJ, Rybina S, Pozniakovsky A, Mayer TU, Walczak CE, Karsenti E, Hyman AA. 2000. Control of microtubule dynamics by the antagonistic activities of XMAP215 and XKCM1 in Xenopus egg extracts. Nat Cell Biol 2:13–19. doi:10.1038/71330

Tran PT, Paoletti A, Chang F. 2004. Imaging green fluorescent protein fusions in living fission yeast cells. Methods 33:220–225. doi:10.1016/j.ymeth.2003.11.017

Varga V, Helenius J, Tanaka K, Hyman AA, Tanaka TU, Howard J. 2006. Yeast kinesin-8 depolymerizes microtubules in a length-dependent manner. Nat Cell Biol 8:957–962. doi:10.1038/ncb1462

Varga V, Leduc C, Bormuth V, Diez S, Howard J. 2009. Kinesin-8 Motors Act Cooperatively to Mediate Length-Dependent Microtubule Depolymerization. Cell 138:1174–1183. doi:10.1016/j.cell.2009.07.032

Vavylonis D, Kovar DR, O’Shaughnessy B, Pollard TD. 2006. Model of formin-associated actin filament elongation. Mol Cell 21:455–466. doi:10.1016/j.molcel.2006.01.016

Vukušić K, Buđa R, Bosilj A, Milas A, Pavin N, Tolić IM. 2017. Microtubule Sliding within the Bridging Fiber Pushes Kinetochore Fibers Apart to Segregate Chromosomes. Dev Cell 43:11–23.e6. doi:10.1016/j.devcel.2017.09.010

Vukušić K, Buđa R, Tolić IM. 2019. Force-generating mechanisms of anaphase in human cells. J Cell Sci 132. doi:10.1242/jcs.231985

Ward JJ, Roque H, Antony C, Nédélec F. 2014. Mechanical design principles of a mitotic spindle. Elife 3:e03398:1–28. doi:10.7554/eLife.03398

Wieczorek M, Bechstedt S, Chaaban S, Brouhard GJ. 2015. Microtubule-associated proteins control the kinetics of microtubule nucleation. Nat Cell Biol 17:907–916. doi:10.1038/ncb3188

Winey M, Mamay CL, O’Toole ET, Mastronarde DN, Thomas H. Giddings J, McDonald KL, McIntosh JR. 1995. Three-Dimensional Ultrastructural Analysis of the Saccharomyces cerevisiae Mitotic Spindle. J Cell Biol 129:1601–1615.

Yamashita A, Sato M, Fujita A, Yamamoto M, Toda T. 2005. The Roles of Fission Yeast Ase1 in Mitotic Cell Division, Meiotic Nuclear Oscillation, and Cytokinesis Checkpoint Signaling. Mol Biol Cell 16:1378–1395. doi:10.1091/mbc.E04

Yu C, Redemann S, Wu H, Kiewisz R, Yeon T. 2019. Central-spindle microtubules are strongly coupled to chromosomes during both anaphase A and anaphase B. Mol Biol Cell 30:2503–2514. doi:10.1091/mbc.E19-01-0074

Yukawa M, Kawakami T, Okazaki M, Kume K, Tang NH, Toda T. 2017. A microtubule polymerase cooperates with the kinesin-6 motor and a microtubule cross-linker to promote bipolar spindle assembly in the absence of kinesin-5 and kinesin-14 in fission yeast. Mol Biol Cell 28:3647–3659. doi:10.1091/mbc.E17-08-0497

Yukawa M, Okazaki M, Teratani Y, Furuta K, Toda T. 2019a. Kinesin-6 Klp9 plays motor-dependent and -independent roles in collaboration with Kinesin-5 Cut7 and the microtubule crosslinker Ase1 in fission yeast. Sci Rep 9:1–15. doi:10.1038/s41598-019-43774-7

Yukawa M, Yamada Y, Toda T. 2019b. Suppressor analysis uncovers that maps and microtubule dynamics balance with the Cut7/Kinesin-5 motor for mitotic spindle assembly in schizosaccharomyces pombe. G3 Genes, Genomes, Genet 9:269–280. doi:10.1534/g3.118.200896

Yukawa M, Yamada Y, Yamauchi T, Toda T. 2018. Two spatially distinct kinesin-14 proteins, Pkl1 and Klp2, generate collaborative inward forces against kinesin-5 Cut7 in S. pombe. J Cell Sci 131:jcs210740. doi:10.1242/jcs.210740

